# Dual cooperation between HSP70 and the 26S proteasome in co-translational protein quality control

**DOI:** 10.1101/2020.07.10.198036

**Authors:** Guiyou Tian, Cheng Hu, Yun Yun, Wensheng Yang, Wolfgang Dubiel, Yabin Cheng, Dieter A. Wolf

## Abstract

Co-translational degradation via the ubiquitin-proteasome system mediates quality control of 15 – 25% of nascent proteins, a proportion that is known to increase dramatically as a result of proteotoxic stress. Whereas the ubiquitylation machinery involved has been characterized, mechanisms coordinating the proteasomal destruction of ribosome-attached nascent proteins remain poorly defined. In pursuit of such mechanisms, we discovered dual cooperation of the HSP70 family member HSPA1 with the 26S proteasome: First, in response to proteotoxic stress, HSPA1 promotes proteasome recruitment to translating 80S ribosomes in a manner independent of nascent chain ubiquitylation. Secondly, HSPA1, in association with its cognate nucleotide exchange factor HSPH1, maintains co-translationally ubiquitylated proteins in a soluble state required for efficient proteasomal degradation. Both mechanisms conspire to confer thermotolerance to cells and to promote the growth of esophageal cancer cells in vitro and in animals. Consistent with these observations, HSPH1 knockout impedes tumor growth in vitro and in animals and correlates with favorable prognosis in digestive tract cancers, thus nominating HSPH1 as a cancer drug target.

**Highlights:**

- Proteotoxic stress causes translational arrest, co-translational protein ubiquitylation, and proteasome recruitment to ribosomes
- Co-translational proteasome recruitment is independent of nascent chain ubiquitylation but is augmented by HSPA1
- HSPA1-HSPH1 disaggregase confers thermotolerance by maintaining the solubility and proteasomal clearance of ubiquitylated proteins
- Low HSPH1 impedes co-translational thermotolerance and tumor growth and correlates with favorable prognosis in various cancers

## Introduction

In warm-blooded species, proteins are continuously exposed to damaging influences that lead to rapid protein denaturation, frequently exacerbated by chronic states of infection, exposure to toxicants, mal- and overnutrition, or inflammation. A defining feature of misfolded proteins is their tendency to aggregate. Such aggregates are implicated in the pathogenesis of a wide range of human disorders, including neurodegeneration and type II diabetes (Amm *et al*, 2014; Buchberger *et al*, 2010; Hartl *et al*, 2011; Ciechanover & Kwon, 2017). On the flip side, the cytotoxicity of protein aggregates is being exploited for cancer therapy, for example by administration of proteasome inhibitors (Chatterjee & Burns, 2017).

The ubiquity and deleterious consequences of protein misfolding necessitates stringent quality control mechanisms. Chief contributor to protein quality control is a large network of chaperones, factors that assist in the folding or disaggregation of proteins (Hartl & Hayer-Hartl, 2009; Kramer *et al*, 2009; Frydman, 2001). If the load of misfolded proteins exceeds the folding capacity of chaperones, denatured proteins or the resulting aggregates are cleared by selective degradation. One of the best studied such pathways is the degradation of proteins that fail to fold correctly in the endoplasmic reticulum (ER) (Stolz & Wolf, 2010). Such terminally misfolded proteins are retro-translocated into the cytoplasm and cleared by ubiquitin-proteasome-dependent ER-associated degradation (ERAD) and presumably autophagy.

Cells also rapidly degrade abnormal proteins that result from errors in translation and/or unsuccessful post-synthetic folding. Co-translational protein quality control is executed by a complex network of chaperones organized in the vicinity of the ribosome (Pechmann *et al*, 2013; Thommen *et al*, 2017; Döring *et al*, 2017; Kramer *et al*, 2018; Joazeiro, 2019). Nascent polypeptide-associated complex (NAC) is a heterodimeric complex that binds in and around the ribosome exit tunnel (Gamerdinger *et al*, 2019). The ribosome-associated complex (RAC) is composed of a specialized HSP70 protein (HSPA14) and a cognate HSP40 J-domain protein (DNAJC2) (Hundley *et al*, 2005; Otto *et al*, 2005; Jaiswal *et al*, 2011). This complex recruits other HSP70 chaperones – Ssb in yeast and HSPA1 in mammalian cells - to ribosome bound nascent peptides (Jaiswal *et al*, 2011). Yeast Ssb chaperones assist co-translational protein folding, facilitate rapid protein synthesis, and support efficient membrane targeting (Willmund *et al*, 2013; Döring *et al*, 2017; Stein *et al*, 2019).

Proteins that cannot attain a stably folded conformation during synthesis are co-translationally degraded by the ubiquitin-proteasome system. Approximately 15% of nascent polypeptides are thought to be ubiquitylated by E3 ligases such as CCR4/Not, Listerin, UBR4/5, and HUWE1 (Bengtson & Joazeiro, 2010; Panasenko, 2014; Wang *et al*, 2013; Yau *et al*, 2017; Duttler *et al*, 2013), followed by degradation by the 26S proteasome (Schubert *et al*, 2000; Turner & Varshavsy, A., 2000). Considering the preferential sensitivity of nascent proteins to stress-induced ubiquitylation and proteasomal degradation (Yau *et al*, 2017; Kim *et al*, 2011; Medicherla & Goldberg, 2008), the fraction of co-translationally degraded proteins may increase under proteotoxic stress conditions.

Whereas it is well established that chaperones and the proteasome are involved in co-translational protein quality control, mechanisms coupling co-translational folding/misfolding with degradation remain poorly defined. Studying the recruitment of HSP70 (HSPA1) and the 26S proteasome to actively translating ribosomes under normal and heat stress conditions, we uncovered distinct but interconnected roles for HSPA1 in co-translational quality control and thermotolerance, functions that appear critical for the survival and growth of esophageal, oral, and other cancers.

## Results

### Co-fractionation of HSPA1 and the 26S proteasome with polysomes shows differential sensitivity to proteotoxic stress

HSPA1 (HSP70) and the 26S proteasome are known to interact with ribosomes (Guerrero *et al*, 2008; Jaiswal *et al*, 2011; Sha *et al*, 2009; Wang *et al*, 2015), suggesting roles in co-translational protein folding and degradation. In line with this possibility, we observed by sucrose density gradient centrifugation of MCF7 cell lysate, that both 26S proteasome subunits (PSMB1, ADRM1) and HSPA1 co-fractionated with polysomes, albeit with different patterns **(Fig. 1A)**. Whereas HSPA1 co-sedimented with both heavy and light polysomes, proteasome subunits were largely restricted to 80S monosomes and light polysomes under standard culture conditions.

**Figure 1.**
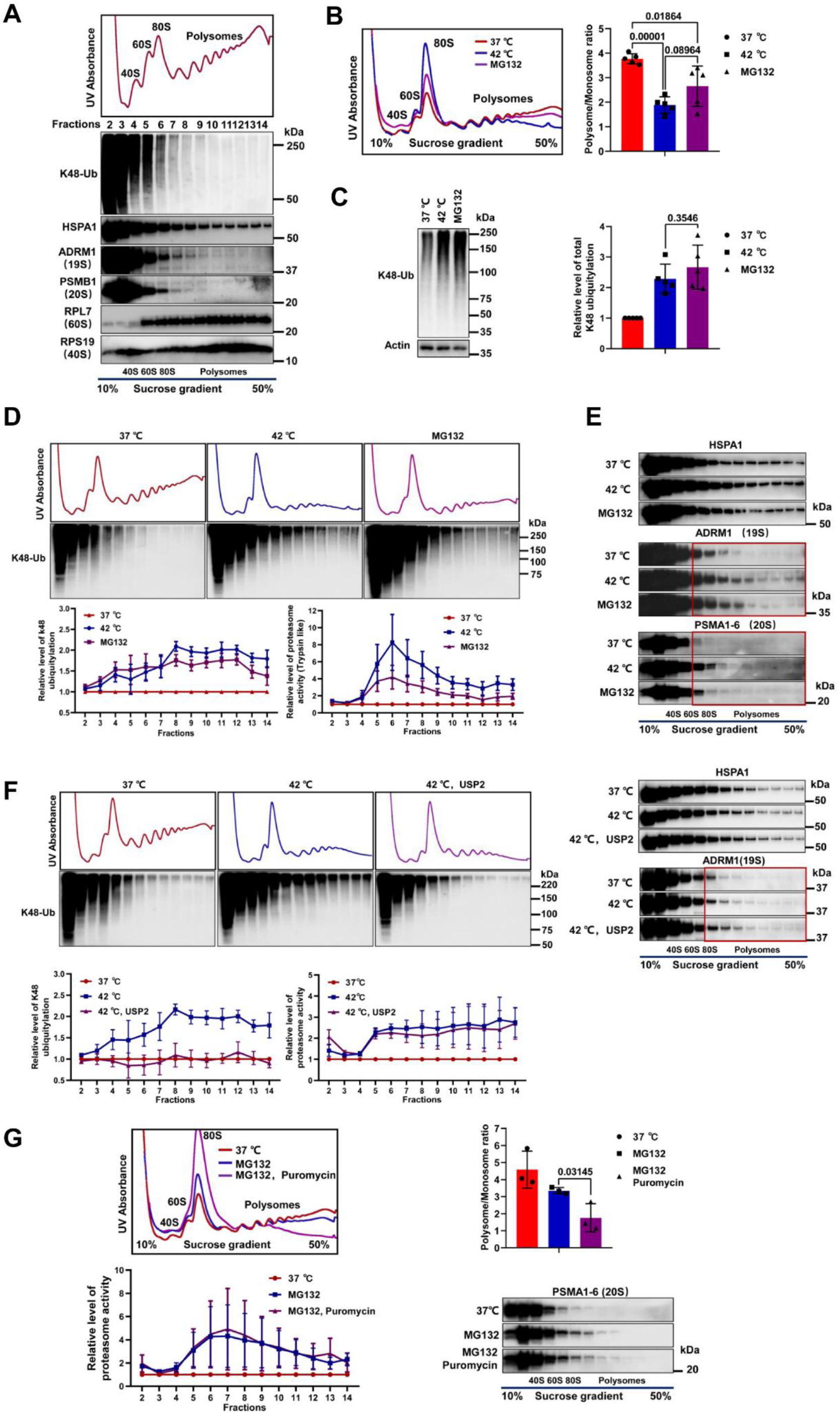
Co-fractionation of HSPA1 and the 26S proteasome with polysomes. **A.** Whole cell lysate of MCF7 cells was fractionated by sucrose density gradient centrifugation and fractions collected along the gradient were assayed for the levels of the indicated proteins by immunoblotting. Absorption at 254 nm was measured across the gradient to trace the elution profile of cellular ribosomes. Monosomes elute in fractions 4 – 6, polysomes in fractions 7– 14 as indicated. **B.** MCF7 cells were exposed to the proteasome inhibitor MG132 (20 μM, 1 h) or to heat shock (42 °C, 1 h) after which an equal amount of cell lysate was subjected to sucrose density gradient centrifugation. Monosomal and polysomal fractions were quantified from the UV traces, and polysome to monosome (P/M) ratios were calculated. Average ratios and the individual datapoints (n = 5) are displayed in a bar graph (means +/− SD). Numbers above the bars represent p values. **C.** MCF7 cells were exposed to MG132 or heat shock as described in B., and total levels of K48-linked polyubiquitylation were determined by immunoblotting and quantified. The bar graph shows average abundance (means +/− SD, n = 5). Numbers above the bars represent p values. **D.** Equal amounts of lysate of MCF7 cells exposed to MG132 and heat shock as described in B. were fractionated by sucrose density gradient centrifugation, and fractions were assayed for the presence of lysine (K) 48-linked polyubiquitylated proteins by immunoblotting with K48 linkage-specific antibodies. The line graph on the left is a quantification of K48-linked polyubiquitin across the gradients relative to the levels obtained in MCF7 cells maintained at 37 °C (means +/− SEM, n = 4). The line graph to the right corresponds to trypsin-like proteasome activity determined in the same fractions using an in vitro assay. Results are shown relative to the proteasome activity obtained in MCF7 cells maintained at 37 °C (means +/− SEM, n = 4). **E.** Equal amounts of lysate of MCF7 cells exposed to MG132 and heat shock as described in B. were fractionated by sucrose density gradient centrifugation, and fractions were assayed for the presence of the indicated proteins by immunoblotting. **F.** Cell lysates were prepared from MCF7 cells maintained at 37 °C or heat shocked at 42 °C for 1h. Where indicated, lysate from heat shocked cells was incubated with recombinant USP2 to remove polyubiquitin chains. Equal amounts of cell lysate were fractionated by sucrose density gradient centrifugation, and K48-linked polyubiquitin was detected by immunoblotting. The line graph on the left is a quantification of K48-linked polyubiquitin across the gradients relative to the levels obtained in MCF7 cells maintained at 37 °C (means +/− SEM, n = 3). The line graph to the right corresponds to trypsin-like proteasome activity determined in the same fractions using an in vitro assay. Results are shown relative to the proteasome activity obtained in MCF7 cells maintained at 37 °C (means +/− SEM, n = 3). Fractions were also assayed by immunoblotting with antibodies detecting the proteasome subunit ADRM1 and the HSP70 chaperone HSPA1 (right panel). **G.** MCF7 cells were exposed to MG132 (20 μM, 1 h) and where indicated, puromycin (100 μg/ml) was added for 10 min prior to harvest and lysate preparation to release nascent peptide chains. Lysates were separated by sucrose density gradient centrifugation (top left), and P/M ratios were calculated from the UV traces (top right, number refers to p value). The elution profile of chymotrypsin-like proteasome activity was measured in gradient fractions and is displayed in a line graph relative to the activity of lysate from cells maintained at 37 °C (bottom left, means +/− SEM, n = 3). The elution profile of 20S proteasome subunits was determined by immunoblotting with an antibody recognizing PSMA1-6 (bottom right).

To determine the polysomal partitioning of HSPA1 and the proteasome upon proteotoxic stress, cells were exposed to elevated temperature (42°C) or to the proteasome inhibitor MG132 (20 μM) for one hour. Neither treatment induced noticeable cell detachment from tissue culture plates or overt cell death (**Fig. EV1A**). Proteotoxic stress inhibited mRNA translation as seen by a decrease in the polysome to monosome ratio (**Fig. 1B**) and caused a marked accumulation of K48 polyubiquitylated proteins (**Fig. 1C**). This accumulation occurred mostly in ribosome-associated fractions of the gradient (**Fig. 1D**), indicating preferential induction of co-translational protein ubiquitylation that is consistent with previous observations demonstrating that nascent proteins are preferentially ubiquitylated and degraded (Duttler *et al*, 2013; Kim *et al*, 2011; Medicherla & Goldberg, 2008; Yau *et al*, 2017). Whereas proteotoxic stress did not notably affect the distribution of HSPA1 along the sucrose gradient, it caused polysomal enrichment of 19S and 20S proteasome subunits (**Fig. 1E**) as well as proteasome activity (**Fig. 1D**), an effect that was more pronounced for heat shock than for MG132. Recruitment of HSPA1 and proteasome subunits to polysomes in response to proteotoxic stress exerted by MG132 was independently confirmed in a different cell line (AD293, **Fig. EV1B**).

Since proteasome recruitment to polysomal fractions appeared to correlate with polysomal ubiquitylation levels (**Fig. 1D**), we asked whether it is due to enhanced ubiquitylation of nascent peptides. To test this, we added recombinant deubiquitylating enzyme USP2 to cell lysate prior to polysome profiling. Despite extensive removal of K48-linked polyubiquitin from polysomal fractions, proteasome recruitment was only minimally affected (**Fig. 1F, Fig. EV1C**). In addition, incubating cell lysate with puromycin prior to sucrose density gradient centrifugation led to the release of ubiquitylated nascent proteins as indicated by a decrease in the polysome to monosome ratio but did not reduce co-fractionation of proteasome subunits or proteasome activity (**Fig. 1G**). These findings suggest that proteasome recruitment to polysomes in response to proteotoxic stress occurs independently of nascent chain ubiquitylation.

### HSPA1 and HSPH1 are selectively induced during recovery from heat shock

Elevated temperature causes the upregulation of heat shock proteins (HSPs), which are known to confer adaptive thermotolerance (Kregel, 2002). To study the role of co-translational protein quality control components in thermotolerance, we rendered MCF7 cells thermotolerant by exposing them to 42 °C for 1 hour, followed by recovery at 37 °C for 4 h (**Fig. 2A**). Throughout this manuscript, MCF7 cells rendered adaptively thermotolerant through this temperature shifting procedure are referred to as MCF7^aTT^ cells. Challenge of heat shock naïve MCF7 cells with exposure to 45 °C led to rapid loss of viability, whereas MCF7^aTT^ cells showed improved survival (**Fig. 2B**). We used this paradigm of adaptive thermotolerance to perform SILAC-based quantitative proteomics of MCF7 cells exposed to 42 °C for increasing times and allowed to recover at 37 °C for various periods. Several chaperones from the HSP10, 20, 60, 70 and 90 families were induced by heat shock for 1 to 4 hours. However, only HSPA1 remained elevated during the recovery (**Fig. 2C**). Remarkably, HSPH1, the cognate nucleotide exchange factor of HSPA1, was not induced during the acute heat shock but only during the recovery (**Fig. 2C**). Selective induction and robust persistence of HSPH1 and HSPA1 during recovery was confirmed by immunoblot analysis, suggesting that these chaperones might mediate thermotolerance. (**Fig. 2D**). As before, heat shock for 1 hour induced the accumulation of K48-linked ubiquitylated proteins in heat shock naïve MCF7 cells but this was not observed in MCF7^aTT^ cells with acquired thermotolerance (**Fig. 2E**). Together, these findings suggest that persistent induction of the HSPA1 and HSPH1, forming a protein disaggregase (Mogk *et al*, 2018; Nillegoda *et al*, 2015; Bracher & Verghese, 2015), mediates adaptive thermotolerance.

**Figure 2.**
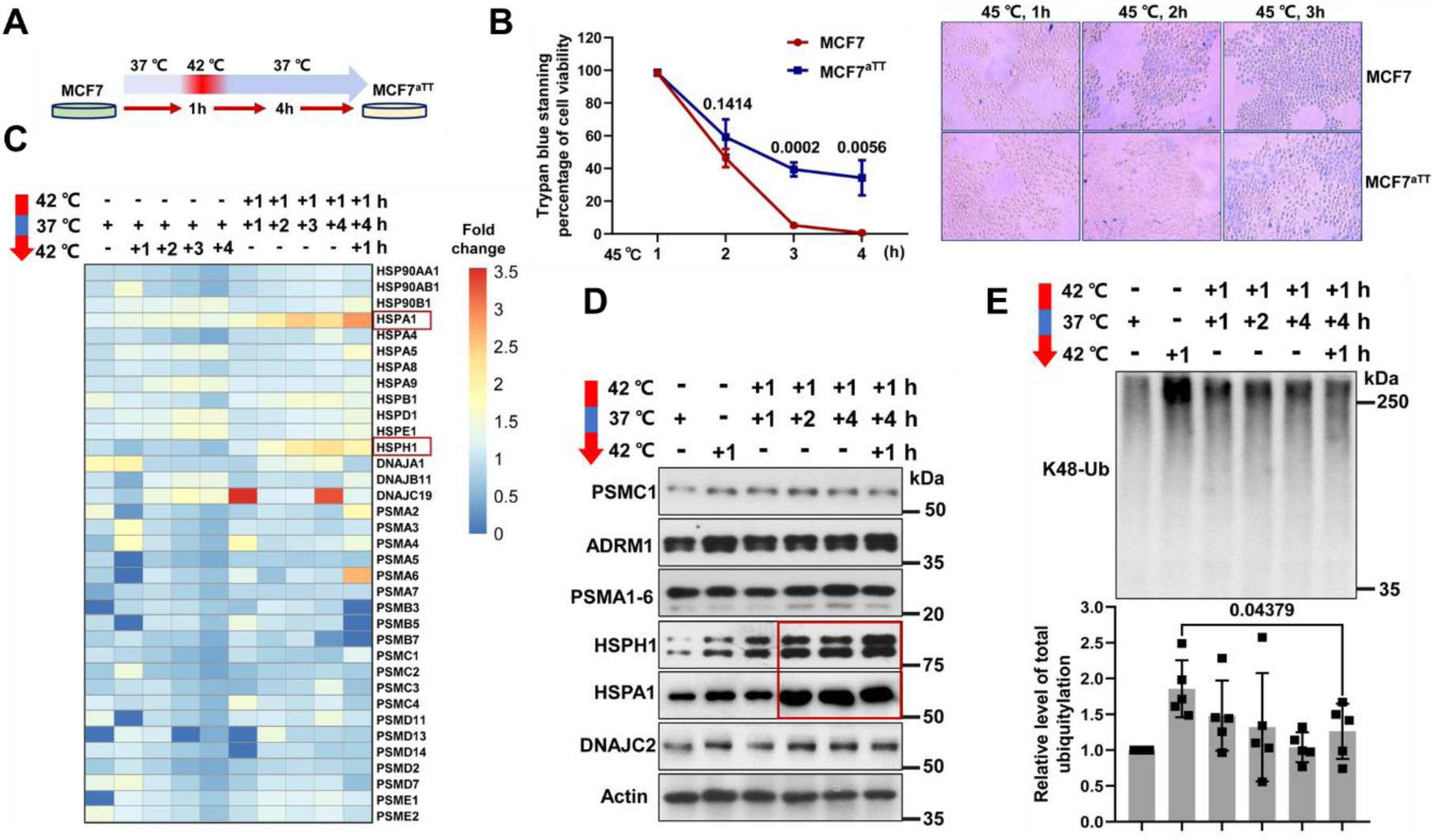
Expression of chaperones during heat shock and recovery leading to thermotolerance. **A.** Schematic diagram of the temperature shift procedure to render MCF7 cells into thermoresistant MCF7^aTT^ cells. **B.** Sensitivity of MCF7 and MCF7^aTT^ cells to heat shock at 45 °C as determined by trypan blue staining. The graph shows means +/− SD, n = 3. **C.** Heatmap showing changes in the expression of chaperones and proteasome subunits in MCF7 cells during heat shock exposure and recovery (MCF7^aTT^) as determined by SILAC proteomics. **D.** The expression levels of the indicated chaperones and proteasome subunits during heat shock and recovery of MCF7/MCF7^aTT^ cells were determined by immunoblotting. **E.** The levels of K48-linked polyubiquitin were determined by immunoblotting of cell lysates prepared from MCF7/MCF7^aTT^ cells kept under the conditions indicated. The bar graph shows a quantification (means +/− SD, n = 3, number indicates p value).

### Adaptive co-translational thermotolerance depends on the activity of HSP70

Nascent peptides are known to be preferential substrates of ubiquitylation in response to heat shock (Michederla & Goldberg 2008). Considering the decrease in bulk cellular ubiquitylation levels induced by heat shock in MCF7^aTT^ cells (**Fig. 2E**), we asked whether co-translational ubiquitylation was also modulated in thermotolerant cells. We performed comparative polysome profiling of naïve MCF7 cells and MCF7^aTT^ cells after exposure to heat shock at 42 °C for 1 hour. Unlike in heat shock naïve cells, renewed heat exposure of thermotolerant MCF7^aTT^ cells resulted in only minor inhibition of protein synthesis and blunted the accumulation of co-translationally polyubiquitylated proteins (**Fig. 3A, B**). This phenomenon, which we refer to as “co-translational thermotolerance”, correlated with a marked increase in the accumulation of HSPA1 in polysomes (**Fig. 3C**).

**Figure 3.**
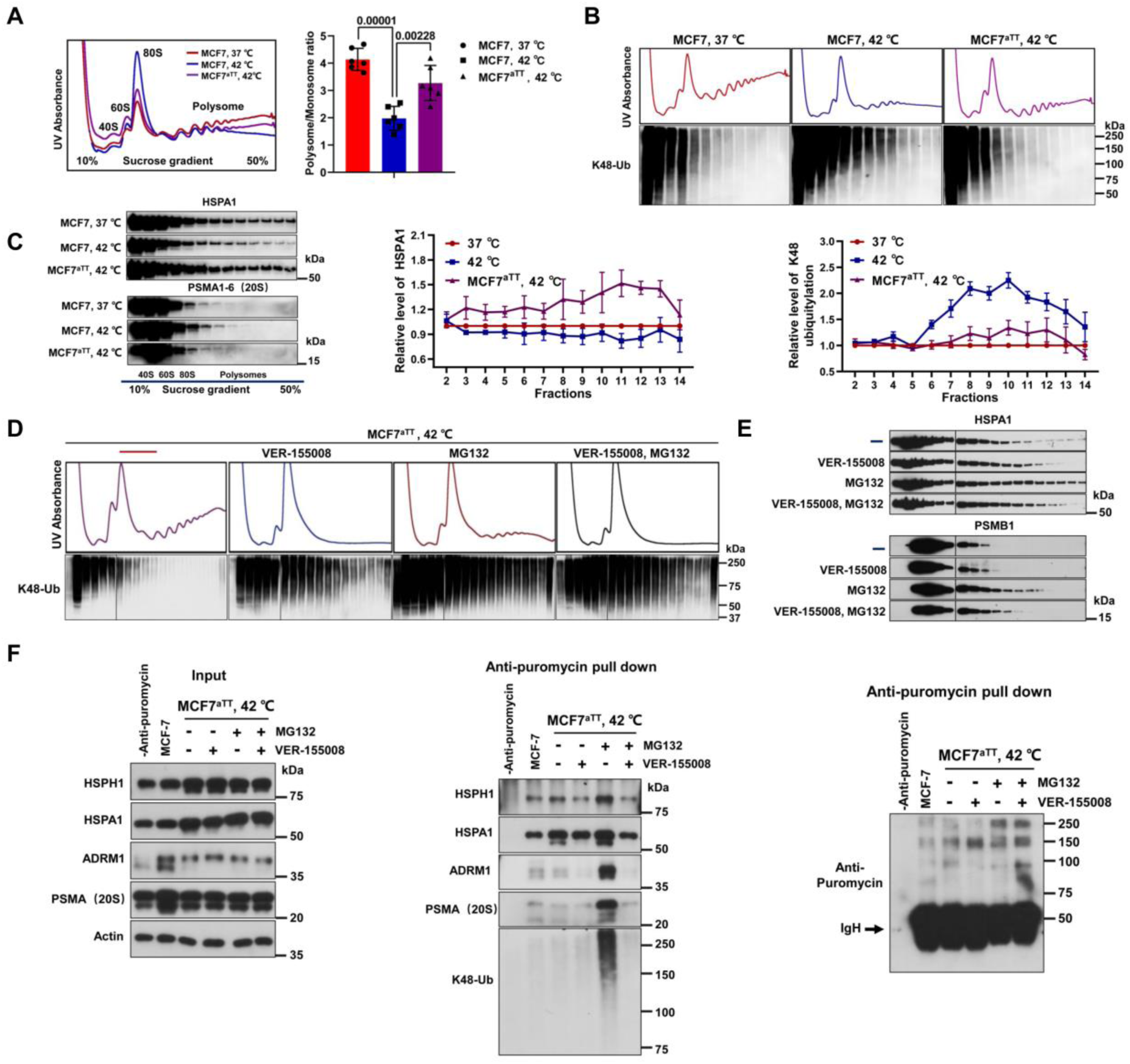
Role of HSPA1 in proteasome recruitment to ribosomes. **A.** Lysate of MCF7 and MCF7^aTT^ cells (rendered thermoresistant through the protocol described in Fig. 2A) were separated by sucrose density gradient centrifugation, and P/M ratios were determined by integration of monosomal and polysomal peak areas (bar graph, mean P/M ratios +/− SD, n = 6). Numbers above bars indicate p values. **B.** Sucrose density gradient fractions described in A. were assayed by immunoblotting for the levels of K48-linked polyubiquitin. Triplicate experiments were quantified with K48-ub levels shown relative to the levels observed in MCF7 cells under standard conditions (37 °C). Data is expressed as means +/− SEM, n = 3. **C.** Sucrose density gradient fractions described in A. were assayed for partitioning of HSPA1 and 20S proteasome subunits by immunoblotting (left panel). Data were quantified in a line graph (right panel, means +/− SEM, n = 4). **D.** MCF7 cells rendered thermoresistant through temperature shifts as described in Fig. 2A (MCF7^aTT^) were incubated with the HSP70 inhibitor VER-155008 (50 μM, 1h) and/or MG132 (20 μM, 1h). Corresponding whole cell lysates were fractionated on sucrose density gradients, and fractions were assayed by immunoblotting for the levels of K48-linked polyubiquitin. Black vertical lines indicate splicing of lanes that were run on different gels due to limitations in lane capacity. **E.** The partitioning of HSPA1 and 20S proteasome subunit PSMB1 along the sucrose gradient described in D. was assessed by immunoblotting. **F.** Total cell lysates were prepared from naïve MCF7 and thermoresistant MCF7^aTT^ cells and assayed by immunoblotting for the levels of the indicated proteins (left panel). The same lysates were employed in immunoprecipitation with anti-puromycin antibodies, and precipitates were assayed for the presence of chaperones and proteasome subunits by immunoblotting (middle panel). The precipitates were also probed with anti-puromycin antibodies to document the pulldown of nascent proteins (right panel).

To determine whether increased HSPA1 recruitment to polysomes mediated co-translational thermotolerance, we exposed MCF7^aTT^ cells to the HSP70 inhibitor VER-155008. Polysome profiling and immunoblotting demonstrated that VER-155008 treatment abolished co-translational thermotolerance as indicated by the re-appearance of K48-linked polyubiquitylated proteins in polysomal fractions after heat shock for 1 hour (**Fig. 3D**). These data suggest that HSPA1 activity is essential for both preventing heat-induced translational arrest and promoting proteasomal degradation of co-translationally ubiquitylated proteins.

Consistent with this conjecture, we found that proteasome inhibition also led to the re-appearance of polysomal polyubiquitylation in heat shocked MCF7^aTT^ cells (**Fig. 3D**). This observation suggests that the decrease in heat-induced co-translational ubiquitylation observed in thermotolerant cells was not a result of decreased modification of nascent peptides with K48 ubiquitin chains but was rather due to increased proteasomal clearance of polyubiquitylated proteins.

Since neither proteasome abundance nor activity and assembly into 26S particles was altered in MCF7^aTT^ cells (**Fig. EV2A,B**), we tested whether decreased heat-induced co-translational ubiquitylation in thermotolerant cells was due to increased proteasome recruitment to polysomes. However, proteasome recruitment to polysomes, as judged by immunoblotting for PSMA1-6 (20S), was slightly decreased rather than increased in heat exposed MCF7^aTT^ cells (**Fig. 3C**). This finding was surprising considering our previous observation that heat shock increased proteasome recruitment to polysomes in naïve MCF7 cells (**Fig. 1E,F**). We therefore explored the possibility that increased recruitment of HSPA1 to polysomes in thermotolerant MCF7^aTT^ cells might promote proteasomal degradation of co-translationally ubiquitylated proteins thus reducing the dwell time of the proteasome on ribosomes. Indeed, when proteasomal degradation was inhibited by MG132, the proteasome appeared trapped in the polysomal fraction of MCF7^aTT^ cells (**Fig. 3E**). This trapping depended on HSPA1 activity as it was reduced by VER-155008 (**Fig. 3E**). The apparent dependence of proteasome recruitment to polysomes on HSPA1 activity was also readily apparent in naïve MCF7 cells exposed to heat shock (**Fig. EV3**).

To biochemically confirm that HSPA1 promotes the recruitment of the 26S proteasome to nascent peptides, we used puromycin labeling and pulldown to isolate nascent peptides from cell lysate of naïve MCF7 cells and MCF7^aTT^ cells before and after heat shock. Some cell samples also received VER-155008 to inactivate HSPA1 or MG132 to inhibit proteasome activity. Binding of proteasome subunits ADRM1 and PSMA1-6 to nascent chains observed in cells treated with MG132 was completely dependent on HSPA1 activity, since it was abolished by VER-155008 (**Fig. 3F**). VER-155008 also reduced the binding of HSPA1 to nascent chains, indicating that catalytic activity might be required for binding. Taken together, these findings suggest an intimate interplay between HSPA1 and the proteasome in co-translational thermotolerance in which HSPA1 promotes the recruitment of the proteasome to heat damaged nascent proteins.

### HSPA1-HSPH1 disaggregase promotes proteasomal degradation by preventing aggregation of ubiquitylated proteins

HSPA1 promotes protein folding and prevents aggregation. Maintaining polypeptide solubility is particularly important to allow for efficient proteasomal degradation (Hjerpe *et al*, 2016; Shiber *et al*, 2013; Wang *et al*, 2011). To study the functional interplay of HSPA1 and the proteasome in the processing and degradation of ubiquitylated proteins induced by proteotoxic stress, we assessed the impact of HSP70 and proteasome inhibitors on the solubility of proteotoxic stress products. Naïve MCF7 and MCF7^aTT^ cells were exposed to heat stress, and the partitioning of ubiquitylated proteins into soluble and insoluble fractions was monitored by immunoblotting. To separate soluble from insoluble proteins, protein aggregates were sedimented by high-speed centrifugation of the cell lysate.

MG132 caused strong induction of polyubiquitin conjugates in MCF7 cells (**Fig. 4A,B**), the majority of which were soluble (**Fig. 4C**). In contrast, whereas heat shock led to an overall lower accumulation of polyubiquitylated proteins, ∼60% of these products were insoluble (**Fig. 4A,B,C**). Incubation of MFC7 cells with HSP70 inhibitor VER-155008 further increased the fraction of insoluble ubiquitylated proteins to ∼80%. In thermotolerant MCF7^aTT^ cells, accumulation of insoluble ubiquitin conjugates in response to heat stress was blunted, and VER-155008 led to only a small increase in insoluble conjugates, most likely due to the high amounts of HSPA1-HSPH1 existing in thermotolerant cells (**Fig. 2B**). Thus, unlike proteasome inhibitor, heat shock and HSP70 inhibition caused accumulation of insoluble ubiquitylated proteins that appear resistant to proteasomal degradation. These data suggest that HSP70 chaperones, and in particular the HSPA1-HSPH1 disaggregase, confer thermotolerance by maintaining the solubility and clearance of proteins that were co-translationally ubiquitylated in response to proteotoxic stress.

**Figure 4.**
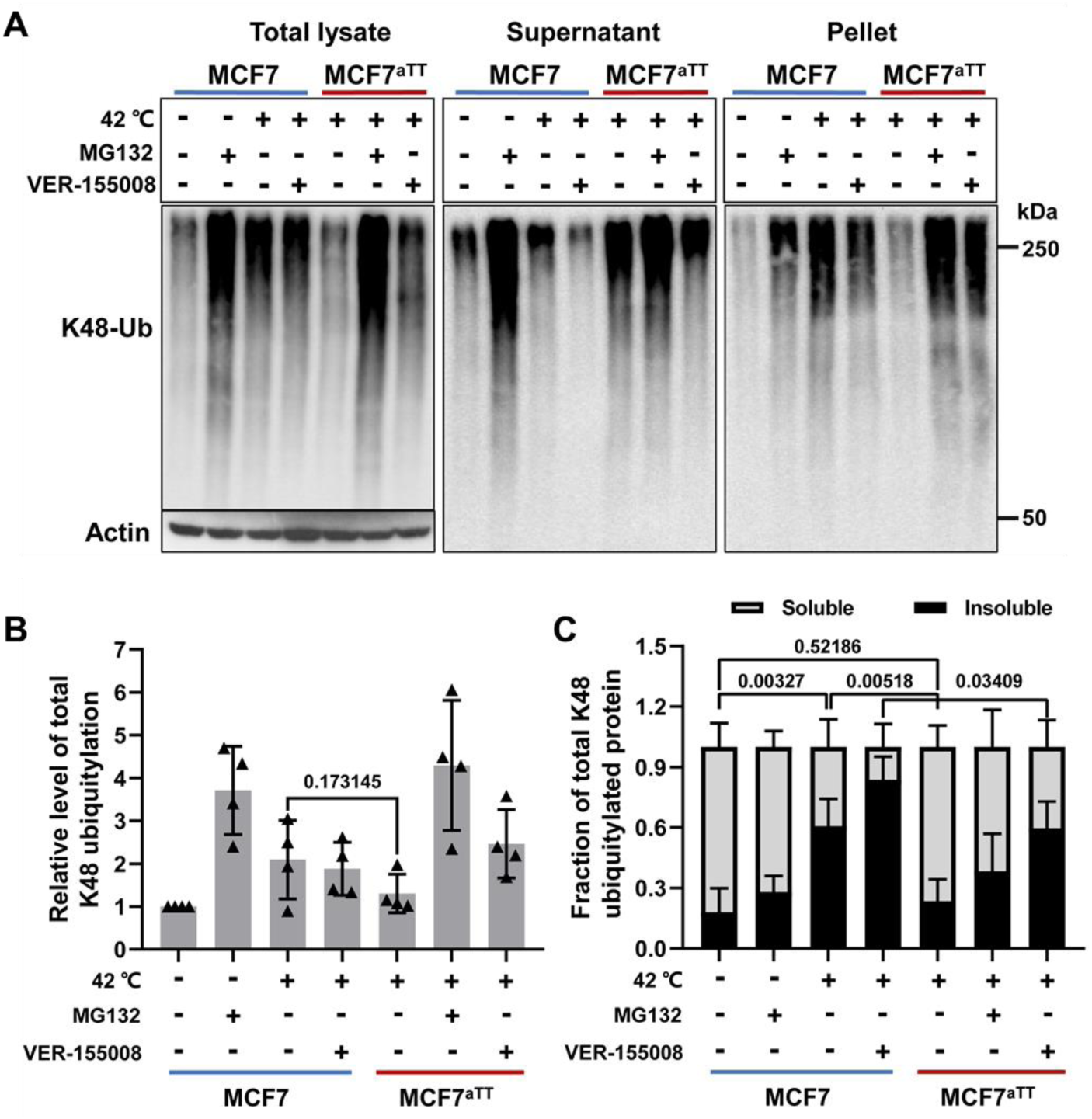
Effect of HSP70 activity on the solubility of ubiquitylated proteins. **A.** Whole lysates of MCF7 and MCF7^aTT^ cells exposed to heat shock and/or proteasome (MG132, 20 μM, 1h) and HSP70 inhibitors (VER-155008, 50 μM, 1h) were fractionated into soluble and insoluble fractions as described in Materials and Methods. Total lysate and fractions were assayed for the level of K48-linked polyubiquitin. Actin is shown as a reference. **B.** The levels of K48-linked polyubiquitin were quantified from multiple independent repeats of the total lysates prepared in A. (means +/− SD, n = 5). **C.** Relative partitioning of K48-linked polyubiquitin into soluble and insoluble fractions was quantified from immunoblots of multiple independent fractionations as shown in A. (means +/− SD, n = 5, numbers above the bars indicate p values).

### Inactivation of HSPH1 curbs thermotolerance of esophageal cancer cells

To test the above conjecture, we sought to genetically inactivate the HSPA1-HSPH1 disaggregase. We were unsuccessful in disrupting HSPH1 by CRISPR/Cas9 in MCF7 cells, possible due to their tetraploid genome. A search of the Cancer Cell Line Encyclopedia revealed esophageal cancer cell lines as the highest expressors of *HSPA1* mRNA among groups of human cancer cell lines derived from 40 different tissues (**Fig. EV4A**). Likewise, esophageal cancer cell lines ranked third highest for expression of *HSPH1* mRNA (**Fig. EV4B**). We therefore attempted to use CRISPR/Cas9 to knock out *HSPA1* and *HSPH1* in the human esophageal cancer cell line KYSE150. Whereas we were unable to disrupt *HSPA1*, we obtained several viable *HSPH1* knockout cell lines (KYSE150^HSPH1-/-^; **Fig. 5A**).

**Figure 5.**
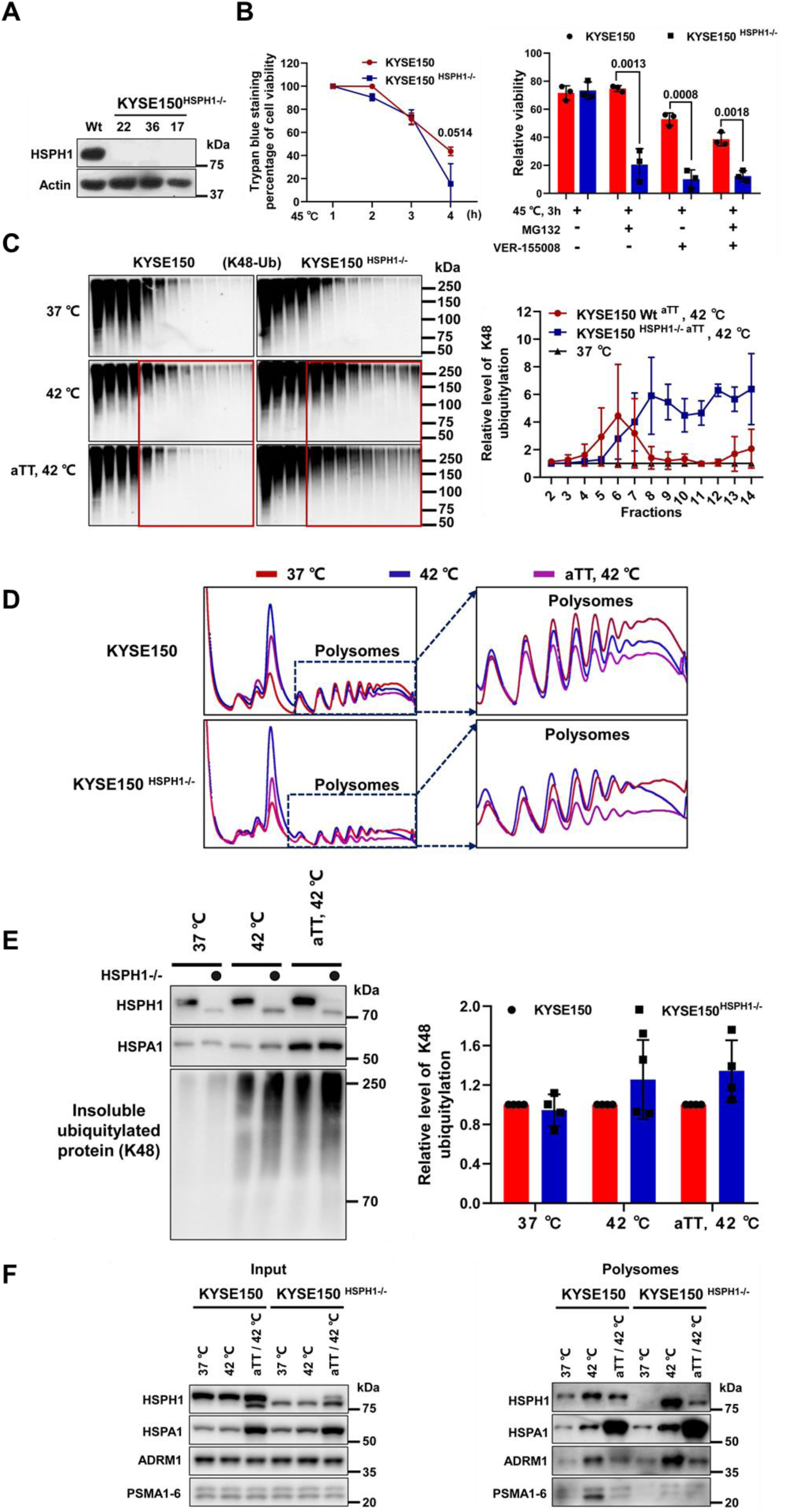
Effect of HSPH1 depletion on co-translational quality control in esophageal cancer cells. **A.** Immunoblot of individual clones of KYSE150 cells in which HSPH1 was disrupted by CRISPR/Cas9 targeting (KYSE150^HSPH1-/-^ cells). **B.** Effect of disrupting HSPH1 on stress sensitivity. KYSE150 and KYSE150^HSPH1-/-^ cells were exposed to heat stress (45 °C, left panel) or proteotoxic stress inducers (right panel), and cell viability was assessed by staining dead cells with trypan blue. Graphs represent means +/− SD, n = 3. **C.** KYSE150 and KYSE150^HSPH1-/-^ cells, either naïve or subjected to the aTT temperature shifting scheme, were exposed to the indicated temperature and cell lysates were separated by sucrose density gradient centrifugation. K48-linked polyubiquitin was detected by immunoblotting of gradient fractions (left panel). Signals were quantified and displayed as a line graph (right panel, means +/− SEM, n = 2). **D.** UV traces of the sucrose density gradients described in C. The polysomal fractions are magnified as indicated. **E.** Whole lysates of cells KYSE150 and KYSE150^HSPH1-/-^ cells exposed to the indicated temperature conditions (aTT scheme and heat shock) were fractionated into soluble and insoluble fractions as described in Materials and Methods. Insoluble fractions were assayed for the level of K48-linked polyubiquitin (left panel). Total levels of HSPA1 and HSPH1 are shown for reference. K48 ubiquitin signals were quantified and displayed in a bar graph (means +/− SDs, n = 4). **F.** Lysates of cells KYSE150 and KYSE150^HSPH1-/-^ cells exposed to the indicated temperature conditions (aTT scheme and heat shock) were obtained and polysomes were isolated by centrifugation through a 30% sucrose cushion. Whole cell lysates (left panel) and the polysomal pellets (right panel) were assayed by immunoblotting for the levels of HSPH1, HSPA1 and proteasome subunits.

KYSE150^HSPH1-/-^ cells were only slightly more sensitive to heat shock at 45 °C than the parental KYSE150 cells, which were relatively heat resistant compared to MCF7 cells (**Fig. 5B**, compare **Fig. 2B**). However, upon additional challenge with MG132 and VER-155008, *HSPH1* knockout cells showed marked thermosensitivity (**Fig. 5B**). KYSE150^HSPH1-/-^ cells were also deficient in developing adaptive co-translational thermotolerance. Whereas parental KYSE150 cells rendered thermoresistant through the same temperature shifting scheme used for MCF7^aTT^ cells (**Fig. 2A**) no longer showed polysomal enrichment of polyubiquitylated proteins after heat shock for 1 hour, KYSE150^HSPH1-/-^ cells subjected to the same treatment still accumulated large amounts of polyubiquitylated proteins in polysomes (**Fig. 5C**). In addition, KYSE150 cells undergoing the adaptive thermotolerance-inducing temperature shift treatment (i.e. KYSE150 ^aTT^ cells) retained a more pronounced translational shut-off in response to heat shock than the corresponding KYSE150^HSPH1-/-^ cells (**Fig. 5D**).

As with VER15008-treated MCF7 cells (**Fig. 4A**), HSPH1-deficient KYSE150 cells were also defective in maintaining the solubility of K48 ubiquitylated proteins accumulating upon heat shock (**Fig. 5E**) thus providing further genetic support for the conception that the HSPA1-HSPH1 disaggregase mediates thermotolerance by solubilizing ubiquitylated proteins in order to enable their clearance by the proteasome.

Heat-induced recruitment of the proteasome to polysomes did not depend on HSPH1 as it occurred in both parental KYSE150 and KYSE150^HSPH-/-^ cells (**Fig. 5F**). This finding indicates that polysomal recruitment of the proteasome uniquely requires HSPA1, either alone or possibly in association with other nucleotide exchange factors such as HSPH2 (aka HSPA4, APG-2). We also observed that KYSE150^aTT^ cells that were rendered thermoresistant and then challenged with heat shock displayed a lower level of proteasome recruitment to polysomes than naïve KYSE150 cells (**Fig. 5F**). This reiterates the observation we previously made in MCF7^aTT^ cells (**Fig. 3C**), which we attributed to a reduced polysomal dwell time of the proteasome in thermoresistant cells based on MG132-induced trapping of the proteasome in polysomal fractions (**Fig. 3E, EV3**). Proteasome recruitment to polysomes was also reduced in KYSE150^HSPH-/-^ cells undergoing the thermotolerance-inducing scheme followed by heat shock (**Fig. 5F**) even though they accumulated large amounts of K48 ubiquitin in polysomes (**Fig. 5C**) which is most likely in an insoluble, proteasome-resistant state (**Fig. 5E**). This observation is consistent with our finding that proteasome recruitment to nascent proteins is largely independent of ubiquitylation (**Fig. 1F**) and suggests an unknown mechanism by which the proteasome disengages from ribosomes when it encounters degradation-resistant ubiquitin conjugates.

### HSPH1 expression is increased in cancer and is required for tumor formation in mice

To put our in vitro observations into a physiological context, we turned to esophageal squamous cell carcinoma (ESCC), the fourth most common cancer in China, which has been firmly linked to intake of food and beverages at high temperature (Tai *et al*, 2017; Whiteman, 2009). We performed immunohistochemistry on paraffin-embedded tissue samples of a set of ESCCs and oral squamous carcinoma (OSCCs) and matching normal tissues. Quantitative scoring showed strong overexpression of cytoplasmic and nuclear HSPH1 in ESCCs and OSCCs compared to the surrounding normal tissue (**Fig. 6A, B**). Whereas no difference was observed between tumors and controls for the expression of HSPA1 in these sets of samples, high *HSPH1* mRNA in a series of 182 ESCCs strongly correlated with poor prognosis (p = 0.035; **Fig. 6C**). Similar correlations with unfavorable prognoses exist for large panels of head and neck and liver cancers (**Fig. EV5A**). These data suggest that cancers associated with intake of hot foods and potentially other cancers obtain a growth and survival advantage by upregulating the co-translational thermotolerance pathway.

**Figure 6.**
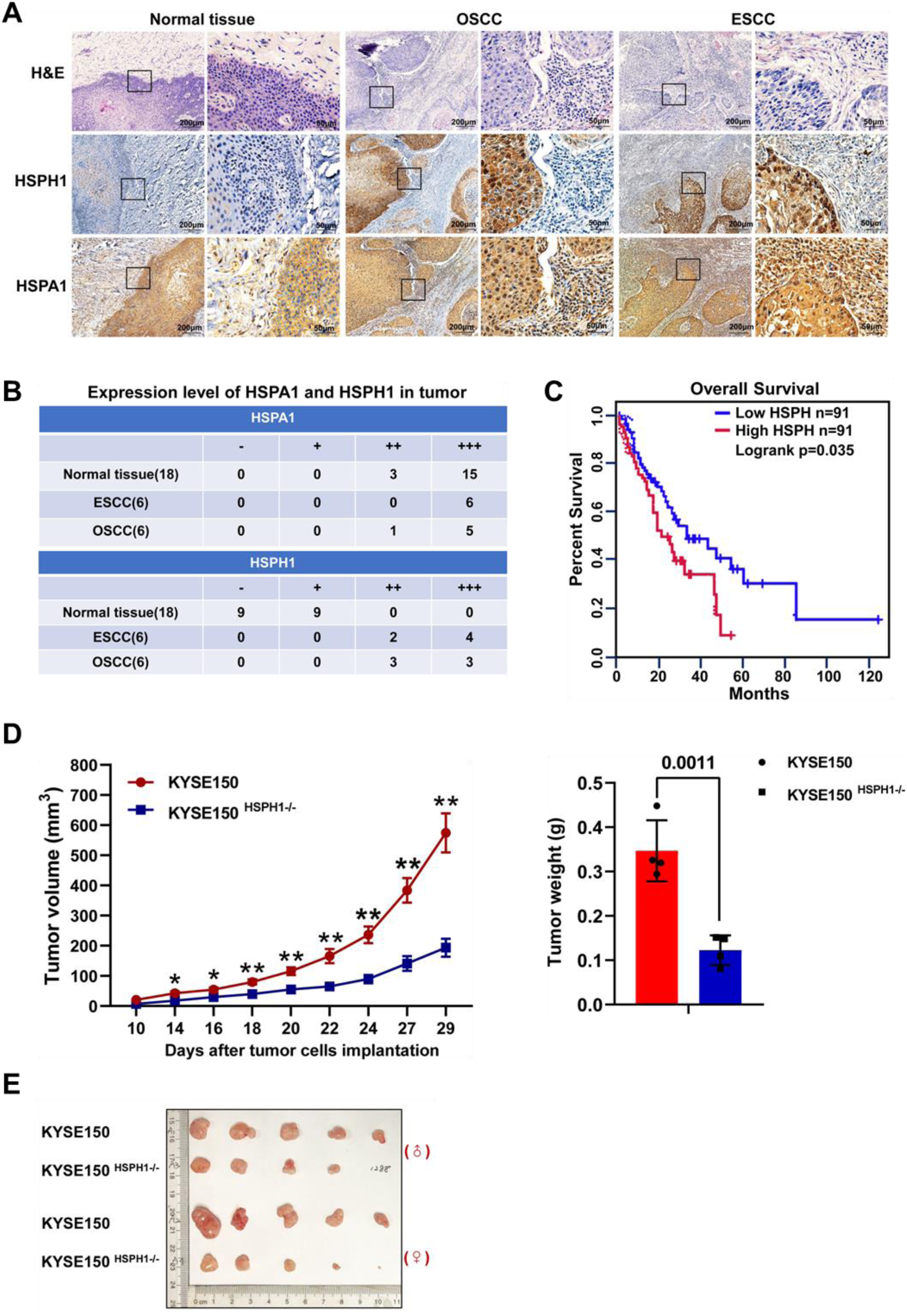
Expression of HSPA1 and HSPH1 in human carcinomas. **A.** Representative micrographs of tissue sections derived from oral squamous cell carcinomas (OSCC) and esophageal squamous cell carcinomas (ESCC) stained with the indicated antibodies by immunohistochemistry (IHC). Top panels show routine H&E staining. Black squares point to regions magnified in the adjacent panels to the right. **B.** IHC staining of HSPA1 and HSPH1 was performed for the indicated numbers of normal, OSCC and ESCC tissues. Staining intensity was scored as described in Materials and Methods. **C.** Kaplan-Meier plots showing the correlation between HSPH1 mRNA expression level and the survival of patients with ESCC. The expression data was obtained from and visualized with KM Plotter (www.kmplot.com, (Nagy *et al*, 2018) using the Pan Cancer algorithm at default settings. **D.** KYSE150 and KYSE150^HSPH1-/-^ cells were injected into nude mice, and xenograft tumor growth was monitored over time (left panel; mean tumor volume +/− SEM, n = 4 with 5 animals per n). Mean tumor volumes at the end of the experiments were determined (+/− SEM, n = 4 with 5 animals per n, number above the bar graph represent p values). **E.** Macroscopic images of representative tumors derived from KYSE150 and KYSE150^HSPH1-/-^ cells (shown are tumors from two of the four independent experiments).

To test this conjecture, we measured the ability of parental KYSE150 and KYSE150^HSPH1-/-^ cells to form tumors in nude mice. Cells were injected subcutaneously, and tumor sizes were measured over a period of 29 days. Tumors derived from KYSE150^HSPH1-/-^ cells were robustly impaired in growth as shown by measurement of average tumor sizes over time, mean terminal tumor weights, and individual tumor sizes (**Fig. 6D, E**). Immunohistochemistry and immunoblotting confirmed severe depletion of HSPH1 protein in KYSE150^HSPH1-/-^ tumors (**Fig, EV5B**). We also ascertained that the differences in tumor sizes were not reflected in different body weights of host animals (**Fig. EV5C**).

## Discussion

Although a substantial fraction of nascent peptides is known to be destroyed co-translationally (Schubert *et al*, 2000; Turner & Varshavsy, A., 2000), mechanisms contributing to co-translational protein degradation remain poorly defined. Unlike with quality control of stalled ribosomes which are split into 40S and 60S subunits prior to ubiquitylation, chaperone-mediated extraction, and proteasomal targeting of the nascent peptides (Joazeiro, 2019), co-translational degradation proceeds while nascent peptides are emanating from elongating 80S ribosomes (Turner & Varshavsy, A., 2000). This necessitates close physical proximity between 80S ribosomes and the 26S proteasome. By determining ribosome recruitment of HSP70 and the proteasome, our study reveals principles of the cooperation between these molecular machines in co-translational degradation and supports a framework mechanistic model of co-translational protein quality control and thermotolerance with implications in oncology (**Fig. 7**).

**Figure 7.**
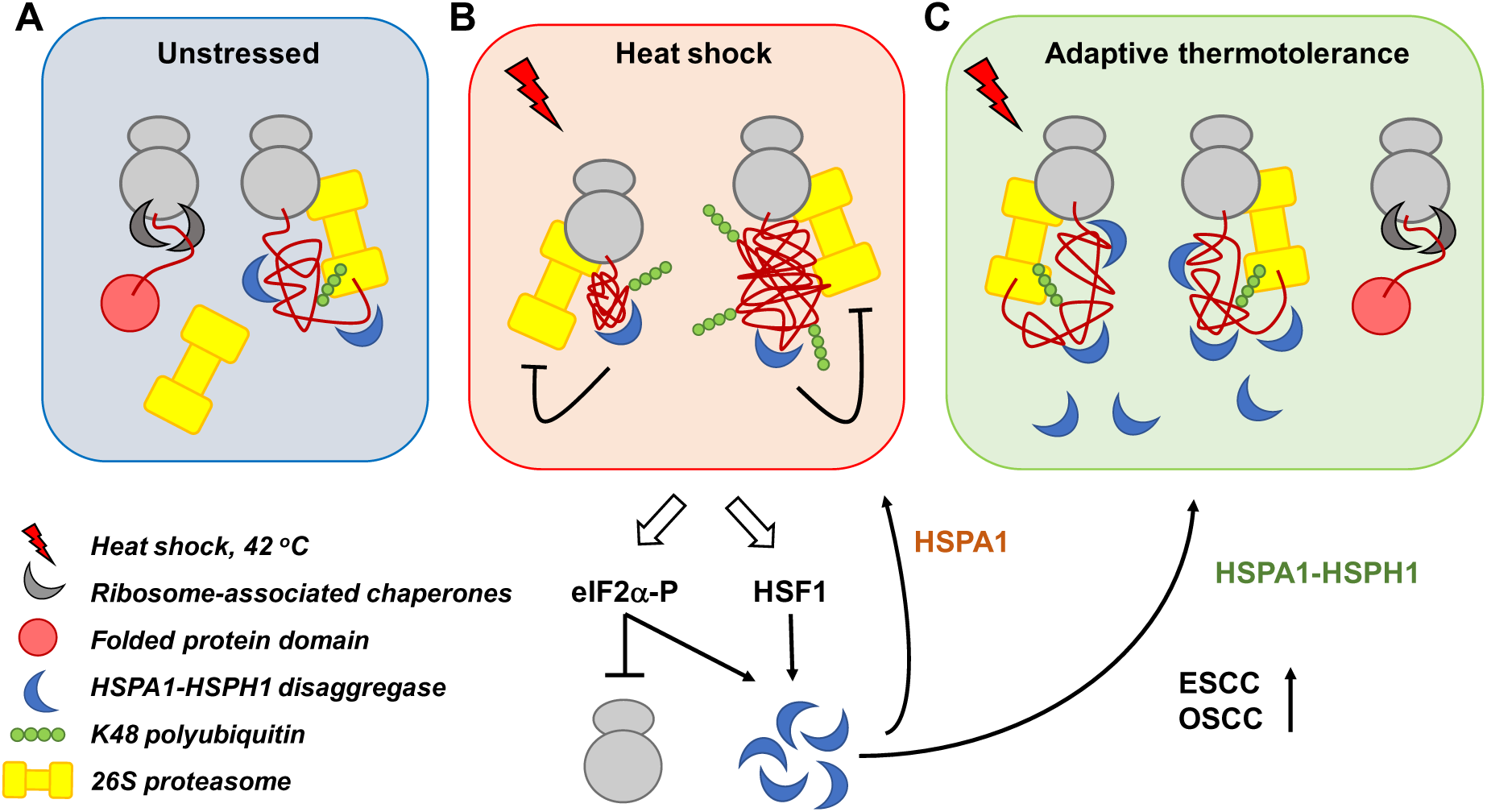
Model of co-translational protein quality control and thermotolerance. (**A**) Under routine conditions, cells co-translationally fold newly synthesized proteins with high efficiency (left panel). A modest fraction of nascent peptides that cannot fold correctly will be ubiquitylated and targeted for co-translational degradation by the proteasome. Proteasomal targeting is aided by the HSPA1/HSPH1, which is present in sufficient amounts to prevent the aggregation of damaged nascent peptides that would impede proteasomal clearance. (**B**) In response to acute heat shock, the HSPA1/HSPH1 become limiting, resulting in the aggregation of ubiquitylated nascent polypeptides. The aggregation prevents clearance by the proteasome. Limitation in HSP70 activity leads to inhibition of global protein synthesis through phosphorylation of eIF2α and mechanisms impinging on translation elongation. Simultaneously, activation of HSF1 leads to transcriptional induction and synthesis of HSPA1. (**C**) In cells rendered thermotolerant through previous heat exposure and recovery, increased levels of the HSPA1-HSPH1 disaggregase facilitates both proteasome recruitment to ribosomes and the solubility of polyubiquitylated nascent chains thus promoting their rapid degradation by the proteasome. As a result, eIF2α phosphorylation subsides, and cells are able to maintain protein synthesis despite the environmental challenge. Tumors, including esophageal and oral squamous carcinomas appear to gain a survival advantage by upregulating this pathway.

### Unstressed condition

We provide evidence that 26S proteasomes associate with actively translating polyribosomes in unstressed cells (**Fig. 7A**). Based on co-fractionation on sucrose density gradients, the association correlates with the presence of K48-linked polyubiquitin. Polysomal polyubiquitylation likely reflects the substantial rate of ubiquitin-proteasome-system (UPS)-dependent degradation of nascent proteins in unstressed cells (Wang *et al*, 2013; Duttler *et al*, 2013; Schubert *et al*, 2000; Turner & Varshavsy, A., 2000). Polysomal proteasome recruitment may, however, not depend on nascent chain ubiquitylation because the proteasome has an intrinsic affinity for 80S ribosomes. This is supported by abundant interactions between proteasome subunits and ribosomal proteins reported in previous studies (Guerrero *et al*, 2008; Sha *et al*, 2009; Wang *et al*, 2015; Verma *et al*, 2000), observations that pointed to an integrated particle referred to as the “translasome” (Sha *et al*, 2009). Based on our published quantitative proteomics data, we estimate that no more than 10% of ribosomes are associated with 26S proteasomes in unstressed yeast cells (Sha *et al*, 2009). This estimate raises the intriguing possibility that the interaction may be dynamically regulated under stress conditions.

### Acute heat stress

Indeed, we found that proteasome recruitment to polysomes is inducible by proteotoxic stress (**Fig. 7B**). Enhanced recruitment coincides with enhanced polysomal polyubiquitylation but does not depend on it. Rather, HSPA1 which is induced during heat shock, is implicated in the recruitment process, since recruitment is suppressed by the HSP70 inhibitor VER155008.

The accumulation of K48-linked polyubiquitin chains in polysomes of stressed cells suggests that co-translational protein degradation is inhibited in the initial response to acute heat stress (**Fig. 7B**). Despite enhanced polyubiquitylation as well as enhanced proteasome recruitment to ribosomes, nascent proteins appear resistant to proteasomal degradation and deubiquitylation. Mechanistically, this may involve inhibition of proteasomal degradation by autoubiquitylation of proteasome subunit ARDM1/RPN13 (Besche *et al*, 2014), although additional mechanisms are possible. A transient period of inhibition of proteasomal degradation may afford chaperones the time needed to refold heat damaged proteins (Wallace *et al*, 2015). At the same time, terminally misfolded nascent proteins may form insoluble aggregates which, despite being marked with ubiquitin, are unsuitable substrates for the proteasome. It is well established that insoluble ubiquitylated proteins are resistant to proteasomal degradation (Shiber *et al*, 2013; Wang *et al*, 2011), unless they are re-solubilized by the HSPA1-HSPH1 disaggregase (Hjerpe *et al*, 2016). We surmise that HSPA1-HSPH1 availability is limiting during the acute stress response. As a direct consequence of exhaustion of HSP70 capacity, protein synthesis is inhibited by (i.) activation of HRI kinase resulting in eIF2α phosphorylation (Lu *et al*, 2001; Thulasiraman *et al*, 1998) and (ii.) through mechanisms that impact translation elongation (Shalgi *et al*, 2013; Liu *et al*, 2013; Sanchez *et al*, 2019). With respect to co-translational quality control, the acute stress situation may resolve, once HSP70 pools are replenished by de novo synthesis downstream of activation of transcription factor HSF1 (Fig. 7B).

### Adaptive thermotolerance

Our quantitative proteomic screen revealed that MCF7 cells that acquired adaptive thermotolerance showed more pronounced upregulation of HSPA1 than cells that underwent acute heat shock (**Fig. 7C**). In addition, HSPH1, its cognate nucleotide exchange factor, was exclusively induced in heat adapted cells thus pointing to an important role for the HSPA1-HSPH1 disaggregase in thermotolerance. Thermotolerant cells no longer respond to the primary insult - heat shock at 42 °C for 1 h - with translational shutdown. This finding suggests that HSP70 activity is no longer limiting in adapted cells thus preventing translational shutdown.

Likewise, thermotolerant cells no longer respond to heat shock with accumulation of polyubiquitylated proteins in polysomes. This is not due to reduced accumulation of polyubiquitylated proteins (and thus reduced protein damage) but a result of more efficient proteasomal removal of ubiquitylated, damaged proteins. As shown by HSP70 inhibition and HSPH1 knockout, the increased degradation efficiency of thermotolerant cells is due to selective induction of HSPA1-HSPH1 during the recovery phase. Thus, our study revealed two important roles of HSPA1 in co-translational protein quality control: (i.) It promotes the recruitment of the proteasome to ribosomes, (ii.) together with HSPH1, it disaggregates ubiquitylated nascent proteins to enable their proteasomal degradation. Our model does not rule out an additional role for HSPA1-HSPH1 in the refolding of heat denatured proteins, which is the preferred strategy of yeast cells to handle heat-induced aggregates (Wallace *et al*, 2015). However, the refolding capacity of mammalian cells appears substantially lower than that of yeast cells (Määttä *et al*, 2020), and in the context of co-translational thermotolerance, re-solubilization and rapid proteasomal degradation may be more relevant.

### Implications for cancer

Our results also assign a crucial role to HSPH1 in facilitating the proliferation of cancer cells in tissue culture and in an animal host. Overexpression of HSPH1 has been found in multiple human cancers, including colon cancer, melanoma, and non-Hodgkin lymphoma (Chatterjee & Burns, 2017), whereas siRNA-mediated knockdown induces apoptosis of colon and gastric cancer cell lines (Hosaka *et al*, 2006). Likewise, we found strong overexpression of HSPH1 in cancers derived from tissues that come into frequent contact with hot beverages and thus potentially heat shock. These observations suggest a scenario in which high levels of HSPH1 would promote increased activity of the HSPH1-HSPA1 disaggregase that confers increased thermal resistance to oral and esophageal cancers. More generally, HSPH1-HSPA1 may allow a diverse set of cancers to adapt to the adverse conditions of the tumor microenvironment which is marked by a mutually reinforcing network of metabolic, genotoxic, proteotoxic stresses (Gorrini *et al*, 2013; Luo *et al*, 2009). This may explain the sensitization of KYSE150^HSPH1-/-^ cells to combinations of stressors, including heat, MG132 and VER155008. Significantly, the proposed role of HSPH1 in stress defense and its selective upregulation in diverse cancers nominate it as an attractive target for therapeutic intervention. Indeed, first attempts at targeting HSPH1 suggest this as a promising avenue for future research (Gozzi *et al*, 2020).

## Structured Methods

### Reagents and Tools Table

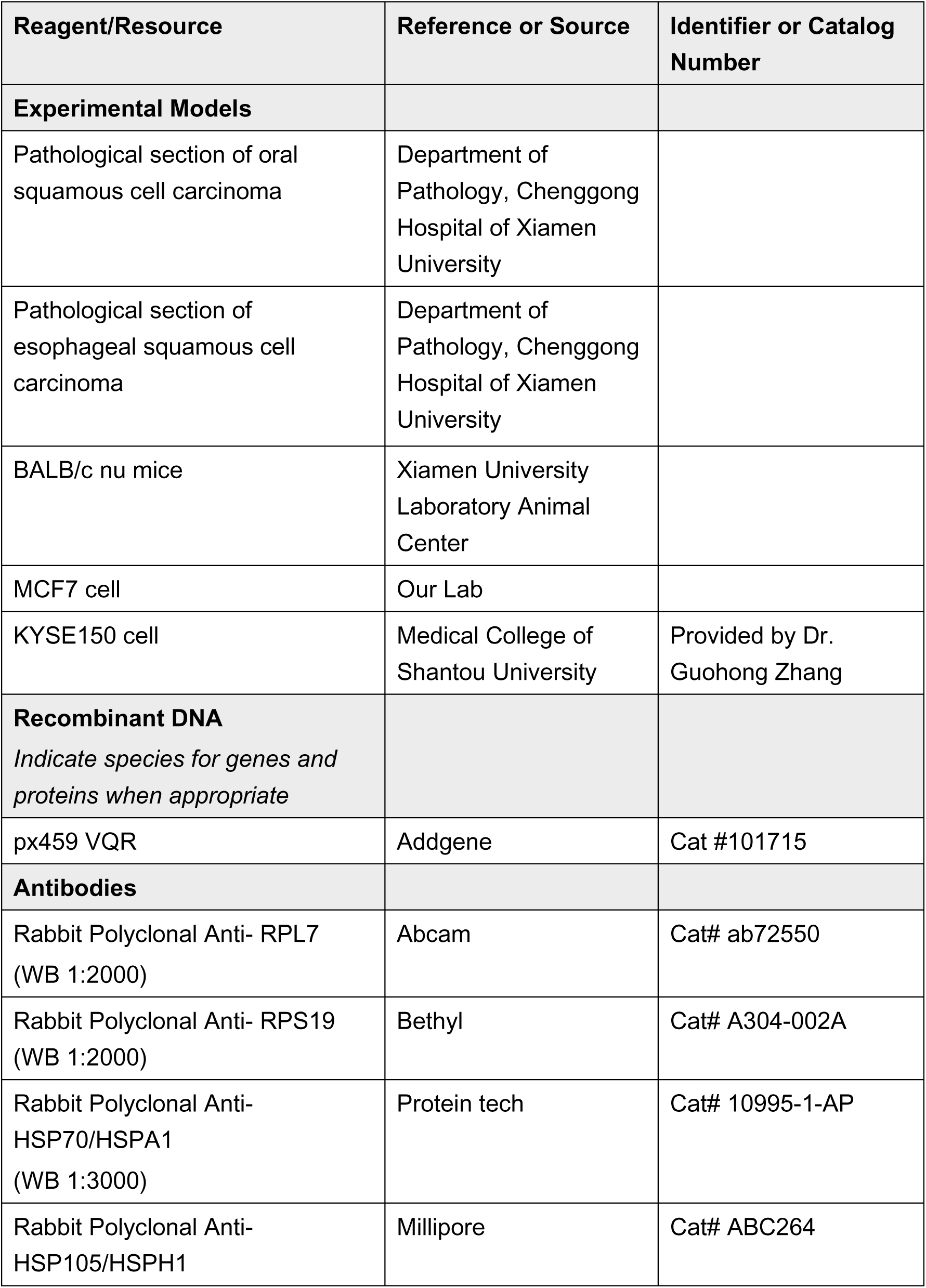
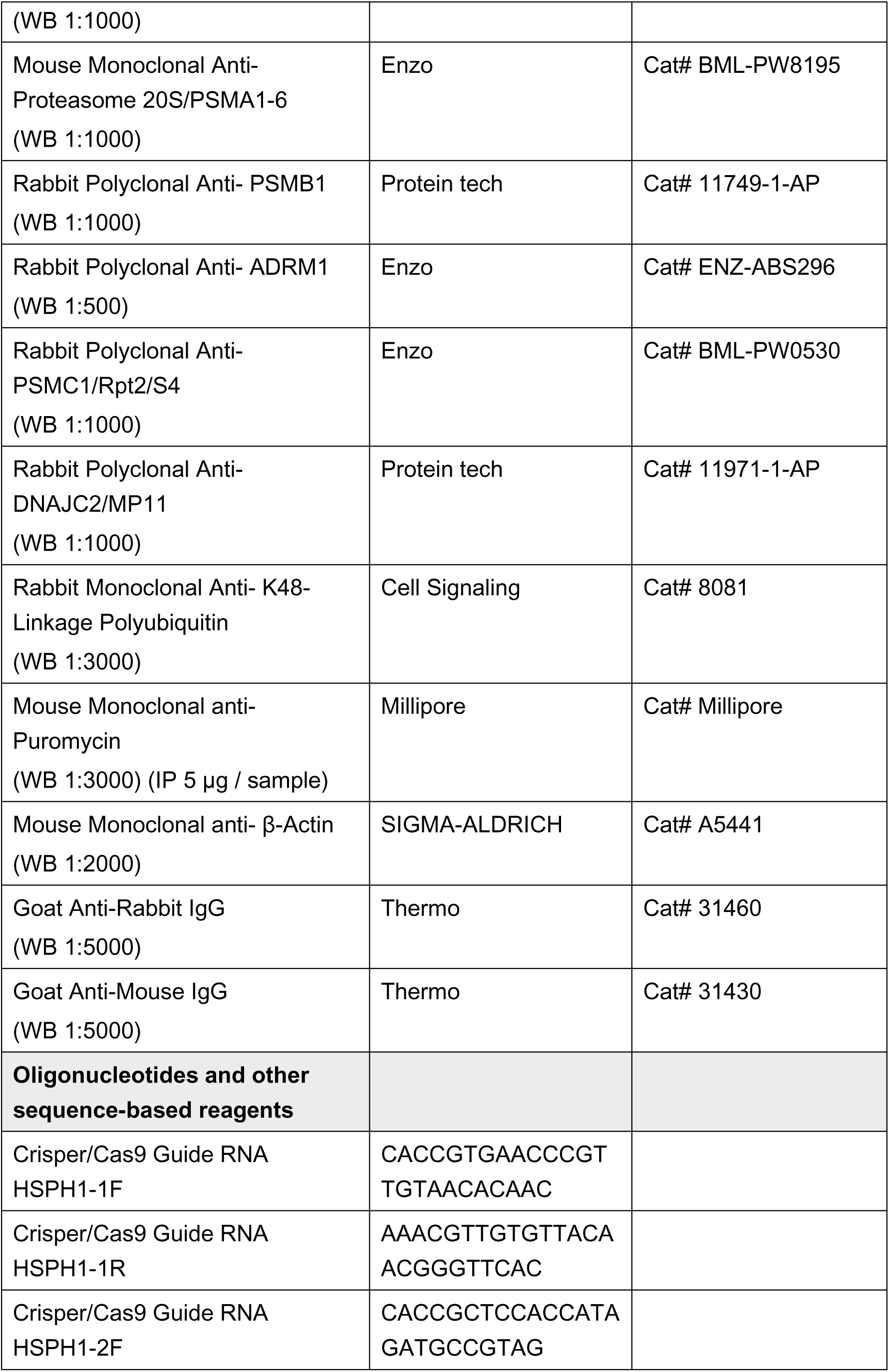
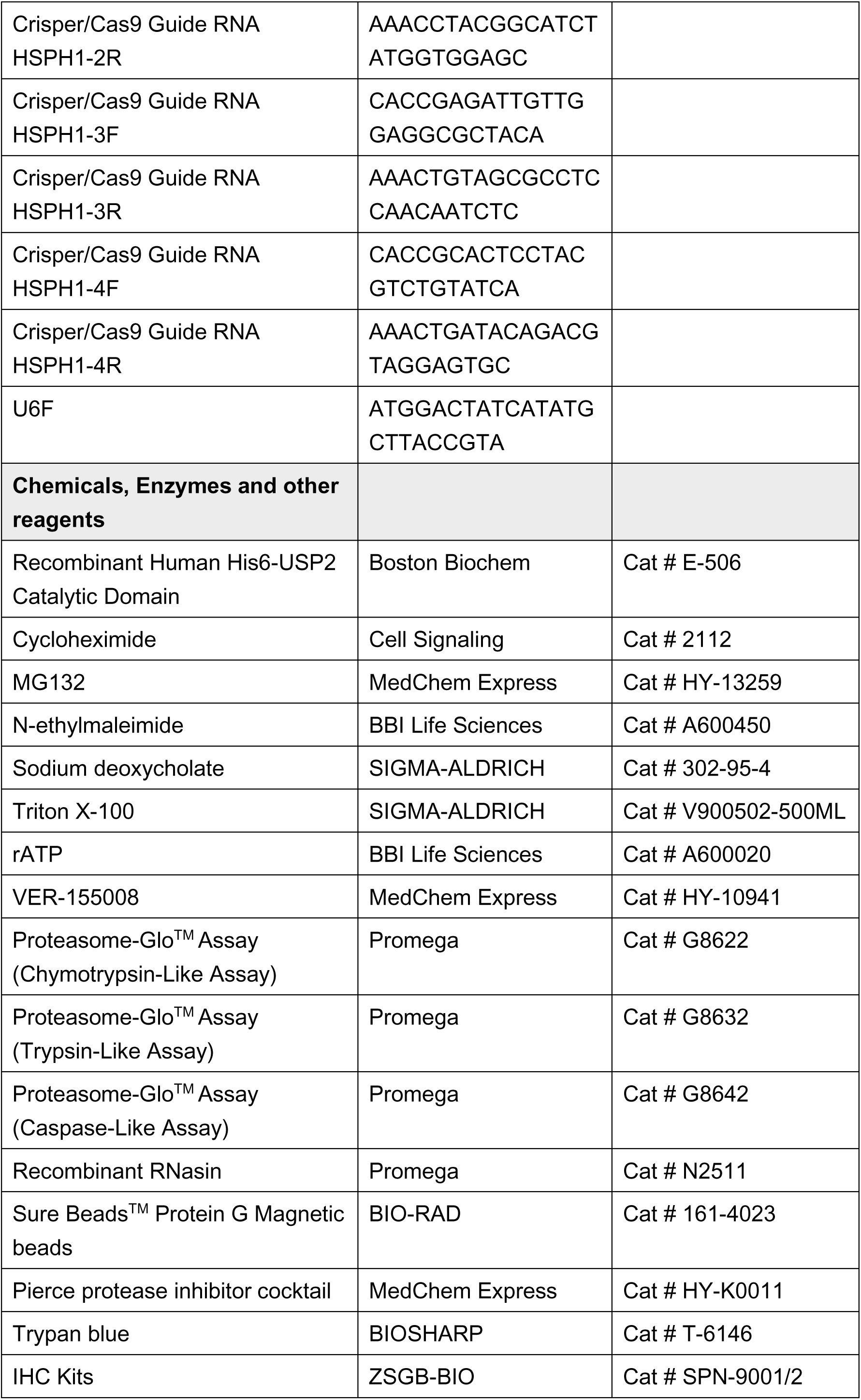
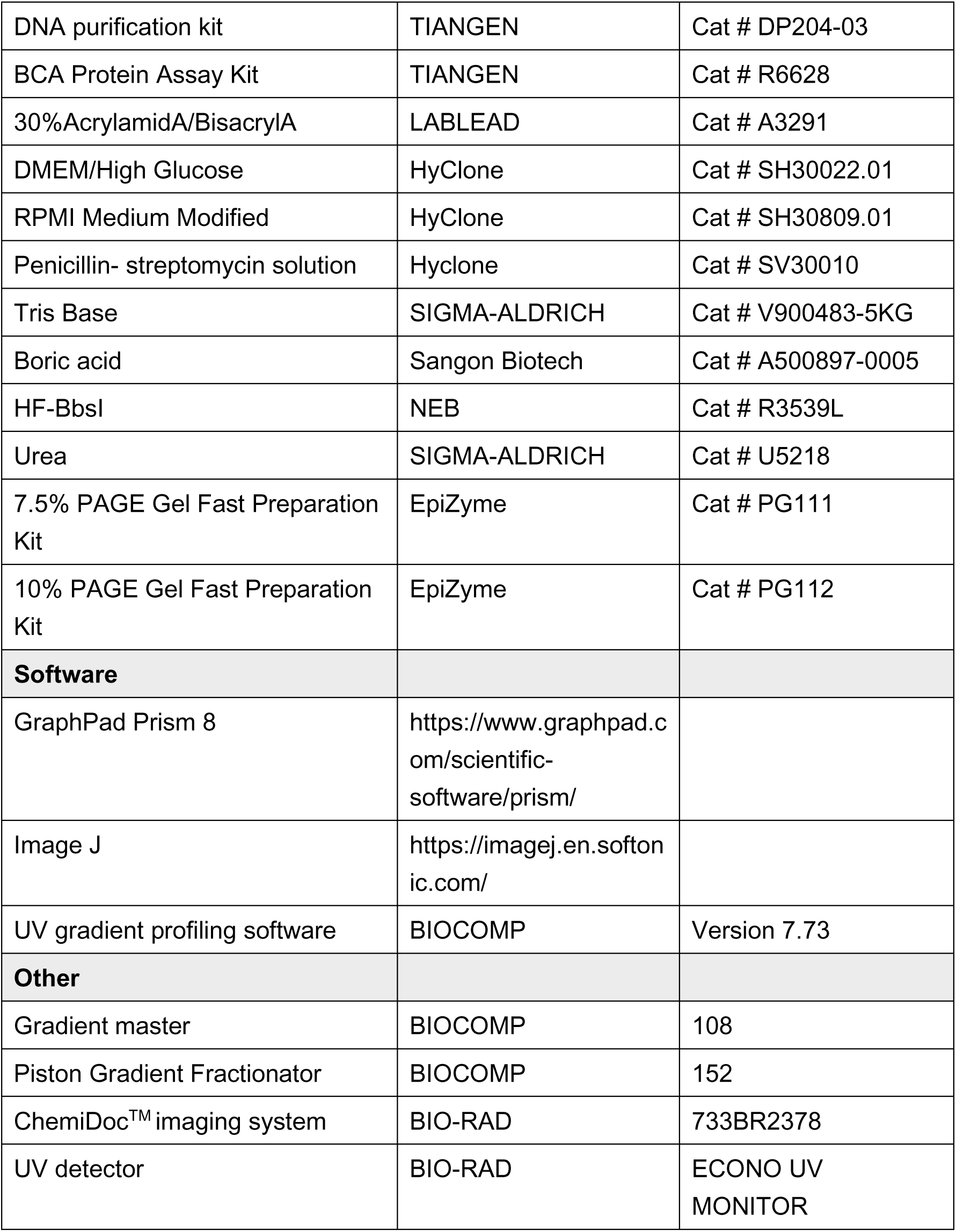

### Cell culture

MCF7 cells were cultured in DMEM/High Glucose (HyClone, SH30022.01), 10% fetal bovine serum (FBS), 100 unit/mL penicillin, and 100 μg/mL streptomycin under 5% CO_2_ at 37°C. KYSE150 cells were cultured in RPMI Medium Modified (HyClone, SH30809.01), 10% fetal bovine serum (FBS), 100 unit/mL penicillin, and 100 μg/mL streptomycin under 5% CO_2_ at 37°C. Cells were authenticated by short tandem repeat sequencing and determined to be free of mycoplasma. For cell viability assays, trypan blue was added to the medium at a final concentration of 0.04% for 2 min. The media was replaced, and cells were qualitatively examined for staining by light microscopy. For quantification of cell viability, cells were trypsinized and counted in a counting chamber. Cell viability was calculated with the formula: %viability = trypan blue stained cells / (unstained cell + stained cells).

### Proteotoxic stress

MCF7 and KYSE150 cells were plated in 6 cm dishes, grown to a density of ∼80% and exposed to heat shock (42°C or 45°C) followed by recovery (37°C) to induce heat stress response and thermotolerance. MG132 (Tocris, 1748) treatments were performed at 20 μM for 1h, VER-155008 (Selleckchem, S7751) treatments were performed at 50 μM for 1h.

### Proteasome activity assay

5 × 10^6^ cells were collected in ice cold PBS. Pellets were resuspended in 0.45 ml hypotonic buffer (5 mM Tris-HCl, pH 7.5, 2.5 mM MgCl_2_, 1.5 mM KCl and 1x Pierce™ protease inhibitor cocktail (MCE, HY-K0011), 1 mM DTT, 2 mM ATP). Cells were incubated in hypotonic buffer on ice for 10 mins and vortexed for 15 seconds. Triton X-100 and sodium deoxycholate were added to final concentration of 0.5% each, and samples were vortexed for another 15 seconds. Cell lysates were centrifuged at 16,500 g for 8 min at 4 °C. Supernatants (cytosolic cell extracts) were collected and proteasome activity was determined followed the descriptions provided with the kit (Promega, G8622).

### Polysome profiling

100 μg/ml cycloheximide (CHX) was added to cells grown in 15 cm dishes 10 mins prior to harvest. 5 × 10^7^ cells were collected in 0.5 ml hypotonic buffer (5 mM Tris-HCl, pH 7.5, 2.5 mM MgCl_2_, 1.5 mM KCl and 1x Pierce™ protease inhibitor cocktail) supplemented with 100 μg/ml CHX (Cell Signaling Technology 2112), 1 mM DTT, and 100 units of RNAse inhibitor (Promega, N2111). Cells were incubated in hypotonic buffer on ice for 10 mins and vortexed for 15 seconds. Triton X-100 and sodium deoxycholate were added to a final concentration of 0.5% each, and the total lysate were vortexed for another 15 seconds. The cell lysates were centrifuged at 16,500 g for 8 min at 4°C. Supernatants (cytosolic cell extracts) were collected and absorbance at OD 260 nm was measured. 20∼30 OD_260_ of lysate was gently layered over 10% – 50% cold sucrose gradients in buffer (20 mM HEPES-KOH, pH 7.4, 5 mM MgCl_2_, 100 mM KCl, 100 μg/mL CHX, 10 units/mL RNAse inhibitor and 1x Pierce™ protease inhibitor cocktail). To prepare the sucrose gradients, 50% sucrose solution (sucrose dissolved in gradient buffer) were layered below a 10% sucrose solution and solutions were mixed using the Gradient Master 108 (Biocomp). Gradients were centrifuged at 36,000 rpm (Beckman, SW41Ti) for 2 h at 4°C. After centrifugation, the polysome profiles were recorded and fractionation through UV detector (BIO-RAD, ECONO UV MONITOR) and the Piston Gradient Fractionator (BIOCOMP, PGF).

For measurement of proteasome activity, 50 μl of each fraction were incubated with proteasome detection reagent (Promega, G8631). For immunoblotting, fractions were mixed with 40 μl 4xSDS sample buffer (200 mM Tris/Cl, pH 6.8, 5% beta-mercaptoethanol, 8% SDS, 40% glycerol, 0.4% bromophenol blue), followed by heating for 10 min at 95°C.

### Digestion of polyubiquitin chains by recombinant USP2

Polysome containing lysate was collected as described above. To digest K48-polyubiquitin chains, the lysate was incubated with 7 μg of recombinant human USP2 catalytic domain protein (Boston Biochem, E-506) for 1 h at 25 °C. After digest, the lysate was layered on top of a 10% – 50% cold sucrose gradient and subjected to density gradient centrifugation.

### Puromycin release of nascent proteins

To release nascent proteins from ribosomes in vivo, 100 μg/ml puromycin was added into culture media for 10 mins prior to collecting cells for lysate preparation and sucrose density gradient centrifugation.

### SILAC sample preparation and quantitative proteomics

MCF7 cells were grown in DMEM containing heavy (^13^C, ^15^N) or light (^12^C, ^14^N) lysine and arginine supplemented with 5% dialyzed FBS for 2 weeks. Cells labeled with heavy amino acids were subjected to the heat shock and recovery treatments indicated in the figures. Control cells were maintained in standard (light) media). Cells were collected and lysed in RIPA buffer (50 mM Tris, 150mM NaCl, 1% NP-40, 0.25% SDS, 1 mM EDTA, and 1 x Pierce™ protease inhibitor cocktail). Protein concentrations were determined by BCA assay, and experimental samples (“heavy”) were mixed with the untreated control sample (“light”) at a ratio of 1:1. 200 μg of protein samples were applied for LC-MS/MS. Filter aided sample preparation (FASP) for LC-MS/MS was done as described (Wiśniewski *et al*, 2009). The digested peptide mixtures were re-dissolved in 0.1% formic acid in ultrapure water prior to LC-MS/MS, and the LC-MS/MS procedures performed as described (Lin *et al*, 2020). Protein identification and quantitation were performed by Thermo Proteome Discoverer (PD 2.1.1.2.) software against UniProt human protein database release 2016_09. Precursor ion mass tolerance was 10 ppm; fragment ion mass tolerance was 0.5 Da. The FDR of protein and peptide was 0.01. The mass spectrometry proteomics data have been deposited to the ProteomeXchange Consortium via the PRIDE partner repository (Perez-Riverol *et al*, 2019) with the dataset identifier PXD020140.

### Separation of soluble and insoluble proteins

5 × 10^6^ cells were subjected to the treatments indicated in the figures, soluble and insoluble protein fractions were prepared by lysing cells in 500 μl hypotonic buffer (supplemented with 20 μM MG132, 50 μM N-ethylmaleimide (NEM), 1 mM DTT, 1 x Pierce™ protease inhibitor cocktail, 2 mM ATP). Triton X-100 and sodium deoxycholate were added to the lysate to a final concentration of 0.5% each. 100 μl of cell lysate was removed (total lysate), and the remainder was subjected to centrifugation at 20,000 x g for 30 min. 120 μl of supernatant (soluble fractions) were mixed with 40 μl 4 X SDS sample buffer, and pellets (insoluble fraction) were dissolved in 120 μl 1 X SDS sample buffer prior to immunoblotting.

### Affinity purification of puromycilated nascent proteins

To label and release nascent peptides by puromycin, 1 × 10^7^ cells were collected and washed with cold PBS, followed by lysis in 500 μl of hypotonic buffer (supplemented with 20 μM MG132, 50 μM NEM, 1 mM DTT, 1 x Pierce™ protease inhibitor cocktail and 2 mM ATP) on ice for 1h with vortexing for 10 s every 10 min. Lysates were clarified by centrifugation at 10,000 rpm for 1 min. 100 μl of the supernatant were removed (total lysate). 100 μg/ml puromycin was added to the remaining lysate, followed by incubation at room temperature for 1h. To pull down puromycilated proteins, anti-puromycin antibody (1:100) was added and incubated at 4 °C for 4h. 50 μl protein-G magnetic beads (BIO-RAD) blocked with 5% BSA and washed in lysis buffer twice were added, followed by overnight incubation. Beads were washed with 500 μl of lysis buffer three times and resuspended in SDS sample buffer.

### Sucrose cushion to enrich polysomal proteins

1 × 10^7^ cells were subjected to the treatments indicated in the figures, and cell lysates were prepared as described for polysome profiling. 120 μl of the lysate were mixed with 40 μl 4 x SDS sample buffer (total sample). 200 μl of cell lysate were layered over a 1 ml cushion of 30% (V/W) sucrose (dissolved in 20 mM HEPES-KOH, pH 7.4, 5 mM MgCl_2_, 100 mM KCl, 100 μg/mL CHX and 1x Pierce™ protease inhibitor cocktail). Samples were centrifuged at 110,000 g (Beckman, TLA-120.2) for 1 h at 4 °C. After centrifugation the pellets were resuspended in 160 μl of 1 x SDS sample buffer and boiled for SDS-PAGE.

### Native PAGE

5 × 10^6^ cells were collected in ice-cold PBS and lysed in 200 μl native lysis buffer (50 mM Tris-HCl, pH 7.4, 1 mM EDTA, 5 mM MgCl_2_, 1 mM DTT, 2 mM ATP) by vortexing with glass beads (3 times for 10 s) at 4 °C. After removal of the glass beads, extracts were cleared by centrifugation at 14,500 g for 10 min at 4 °C. Protein concentration was measured by monitoring OD_280_, and mixed with 1/5 volume of 5 x native loading buffer (250 mM Tris-HCl, pH 6.8, 50% Glycerol, 0.05% bromophenol blue). 20 μg of each lysate were subjected to 4% native PAGE. Native gels were prepared as follows (recipe for one minigel): 2 ml 5 x native buffer (0.45 M Tris base, 0.45 M boric acid, 10 mM MgCl_2_, pH8.1 - 8.4), 1.34 ml 30% acrylamid/bisacrylamide, 0.5 ml glycerol, 6.08 ml H_2_O, 1mM ATP, 1 mM DTT, 80 μl 10% APS, 8 μl TEMED). Gels were left to polymerize at room temperature for 40 min. Gels were run at 75 V for 1h and then at 125 V for 2h in an ice bucket with pre-cold running buffer (0.09 M Tris base, 0.09 M boric acid, 2 mM MgCl_2_, 1 mM ATP, 1 mM DTT).

### Immunoblotting

For immunoblotting, samples were separated by SDS-PAGE or native PAGE and proteins were transferred onto a PVDF membrane. Membranes were blocked in 5% skim milk in TBST for 1h at room temperature. Membranes were probed with primary and secondary antibodies as follows: RPL7 (Abcam, ab72550) 1:2000, RPS19 (Bethyl, A304-002A) 1:2000, Proteasome 20S/PSMA (Enzo, BML-PW8195) 1:1000, ADRM1/RPN13 (Enzo, ENZ-ABS296) 1:500, PSMB1 (Proteintech, 11749-1-AP) 1:1000, PSMC1/Rpt2/S4 (Enzo, BML-PW0530) 1:1000, DNAJC2/MPP11 (Proteintech,11971-1-AP) 1:1000, Anti-Puromycin (Millipore, MABE343) 1:3000, Hsp70/HSPA1 (Proteintech, 10995-1-AP) 1:3000, HSPH1(Millipore-ABC264) 1:1000, K48-Linkage Specific Polyubiquitin (Cell Signaling, 8081) 1:3000, Anti-β-Actin (SIGMA-ALDRICH, A5441) 1:2000, Goat Anti-Mouse IgG (Thermo, 31430) 1:5000, Goat Anti-Rabbit IgG (Thermo,31460) 1:5000. All primary antibodies were diluted in 1% BSA in TBST, and blots were incubated overnight at 4 °C. Blots were washed with TBST (3 times for 10 min). All secondary antibodies were diluted in 5% skim milk in TBST and blots were incubated at room temperature for 1h, followed by washing in TBST (3 times for 10 min). Chemiluminescent detection reagent (Advansta, 191026-09) was added, and chemiluminescence signals were acquired by exposure toX-ray film. Blots that were used for quantifications were visualized with the ChemiDoc^TM^ imaging system (BIO-RAD) in an exposure time accumulation approach, and the appropriate exposure signal were selected for quantification. Quantifications were performed using ImageJ software.

### Immunohistochemistry

Tissue sections from the archives of the Department of Pathology at the Chenggong Hospital of Xiamen University were stained with monoclonal antibodies (HSPH1, 1:50 for 2 h at room temperature; HSPA1, 1:200 for 2 h at room temperature) using the IHC kits (ZSGB-BIO, SPN-9001/2) according to the procedures provided by the manufacturer (ZSGB-BIO). Slides were scored by microscopy, according the staining result the protein expression levels were divided into negative (-, no staining of any tumor cells), weakly positive (+, faint or focal staining), moderately positive (++, strong staining in a minority of cells) and strongly positive (+++, strong signal in the majority of cells).

### Knockout of HSPH1 using CRISPR-Cas9

To establish HSPH1 knock-out cell lines using the CRISPR-Cas9 system, guide RNAs (gRNAs) were designed using a web-based toll (http://crispr.mit.edu/). The vector plasmid px459 VQR was obtained from Addgene (#62988). Vector digest system: 1 μg px459 VQR was digested with HF-BbsI (NEB) and purified with a DNA purification kit (TIANGEN, DP204-03). The corresponding sense and antisense oligonucleotides listed below were annealed by incubation at 37 °C for 30 min, heating to 95 °C for 5 min and cooling to 25 °C at a rate of 5 °C /min. Annealed oligos were ligated into px459 VQR and cloning products were confirmed by sequencing. To induce HSPH1 gene deletion, the plasmids were transfected into KYSE150 cells using Lipofectamine 2000 transfection reagent (Invitrogen) according to the procedures provided by the manufacturer. 48h after transfection, cells were plated for the collection of individual clones. After 2 weeks, cell clones were tested for deletion of HSPH1 by immunoblotting.

gRNA used in this study:

oligo HSPH1-1F: CACCGTGAACCCGTTGTAACACAAC
oligo HSPH1-1R: AAACGTTGTGTTACAACGGGTTCAC
oligo HSPH1-2F: CACCGCTCCACCATAGATGCCGTAG
oligo HSPH1-2R: AAACCTACGGCATCTATGGTGGAGC
oligo HSPH1-3F: CACCGAGATTGTTGGAGGCGCTACA
oligo HSPH1-3R: AAACTGTAGCGCCTCCAACAATCTC
oligo HSPH1-4F: CACCGCACTCCTACGTCTGTATCA
oligo HSPH1-4R: AAACTGATACAGACGTAGGAGTGC

### Tumor xenograft studies

BALB/c nude mice (5–6 weeks of age) were used for subcutaneous xenografts. Mice were maintained in a pathogen-free environment in a 12 h light/dark cycle at 40–60% humidity and 22 – 24 °C with sterilized food and tap water ad libitum. 2.5 × 10^5^ cells in a volume of 100 µl were inoculated subcutaneously into nude mice. After tumor reached a diameter of ∼3mm, tumor sizes were monitored three times per week by measurement with a caliper and body weights were recorded. Tumor volumes were calculated according to the formula: volume = width^2^ × length/2. Mice were sacrificed after three weeks and tumors were excised for analysis. For protein extraction, 100 μl tissue lysis buffer (50 mM Tris, 150 mM NaCl, 0.5 mM EDTA, 1 mM DTT, 1% Triton X-100, 0.5% sodium deoxycholate, 0.1% SDS and 1 mM PSMF) was added to 10 mg tumor tissue, followed by disruption with glass beads (3 times for 10 s) at 4 °C. Extracts were cleared by centrifugation at 12,000 g for 10 min at 4 °C. 20 μl of the supernatant was diluted in 100 μl lysis buffer and 40 μl 4 x SDS sample buffer for immunoblotting.

### Quantification and statistical analysis

All p values were determined using the Multiple t-tests function in GraphPad Prism 8 (two-stage linear step-up procedure of Benjamini, Krieger and Yekutieli with Q set to 5%. Each row was analyzed individually, without assuming a consistent standard deviation).

### Data availability

The mass spectrometry proteomics data have been deposited to the ProteomeXchange Consortium via the PRIDE partner repository (Perez-Riverol *et al*, 2019) with the dataset identifier PXD020140.

## Acknowledgements

This work was funded through grant grants 81773771 and 31770813 from the National Science Foundation of China (D.A.W and W.D.) and from the Fujian Provincial Department of Science & Technology (2017J05138) and the Innovation Program of Xiamen University Department of Life Sciences & Human Health (Y.C). The support from the Equipment Platform of the State Key Lab of Cellular Stress Biology at Xiamen University is gratefully acknowledged.

## Author contributions

Conceptualization, G.T. and D.A.W.; Methodology, G.T., C.H., Y.Y. and W.Y.; Formal Analysis, G.T., Y.C. and W.Y.; Investigation, G.T., Y.Y., C.H. and W.Y.; Writing – Original Draft, G.T and D.A.W.; Writing – Review & Editing, G.T., Y.Y., C.H., W.Y., W.D., Y.C. and D.A.W.; Visualization, G.T., D.A.W.; Supervision, W.D., Y.C. and D.A.W.; Funding Acquisition, W.D., Y.C. and D.A.W.

## Expanded View Figures

**Fig. EV1.**
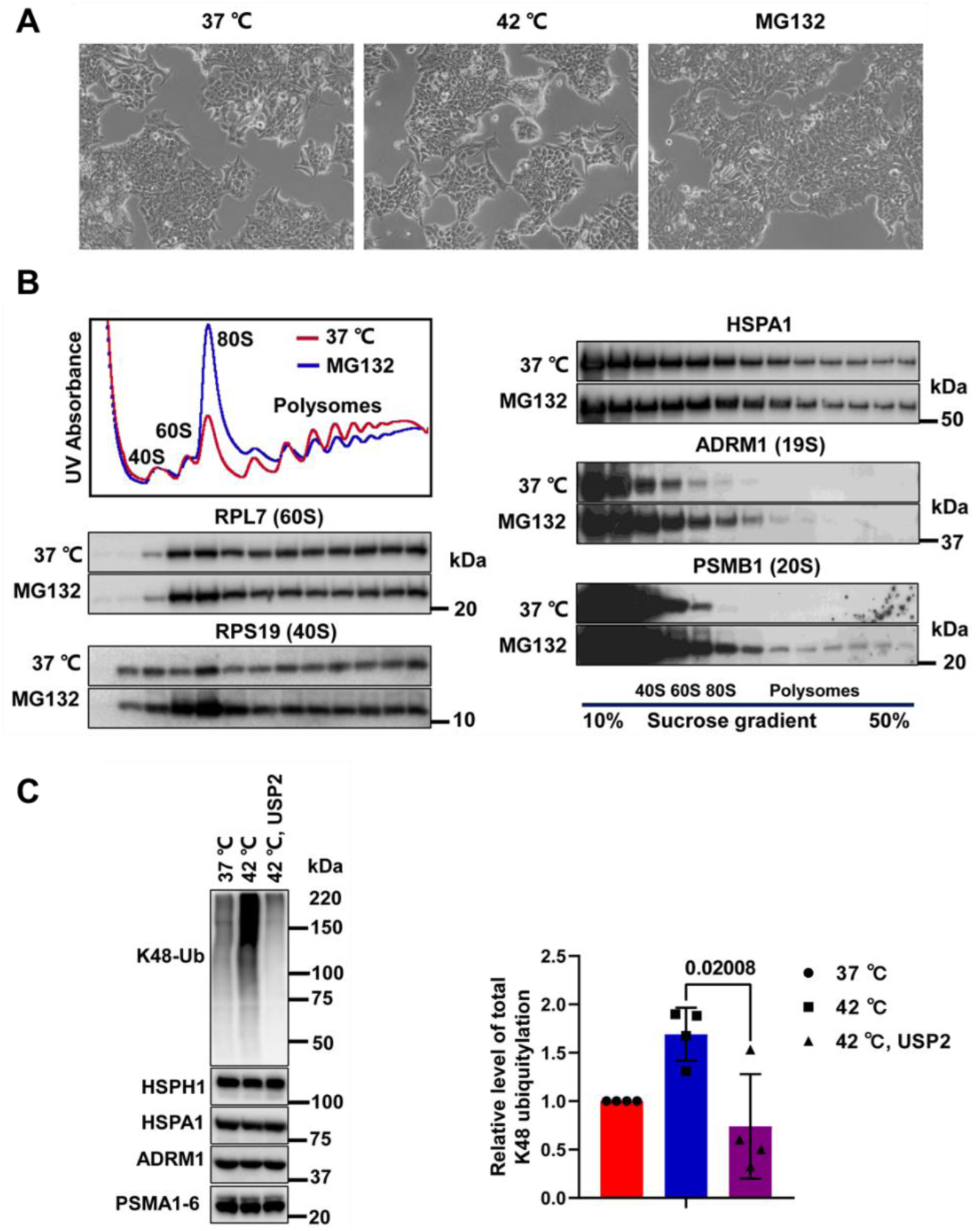
Effect of heat shock on MCF7 cell viability and ubiquitylation levels. **A.** Micrographs of MCF7 cells exposed to the indicated conditions for 1 h. **B.** AD293 cells were exposed to the proteasome inhibitor MG132 (20 μM, 1 h) after which an equal amount of cell lysate was subjected to sucrose density gradient centrifugation (UV traces, top left). The abundance of the indicated ribosomal proteins, chaperones, and proteasome subunits was determined by immunoblotting. **C.** Cell lysates were prepared from MCF7 cells maintained at 37 °C or heat shocked at 42 °C for 1h. Where indicated, lysate from heat shocked cells was incubated with recombinant USP2 to remove polyubiquitin chains. Equal amounts of cell lysate were assayed for the levels of the indicated proteins by immunoblotting. A quantification of the levels of K48-linked polyubiquitin is shown in the bar graph (means +/− SD, n = 4, number indicates p value).

**Fig EV2.**
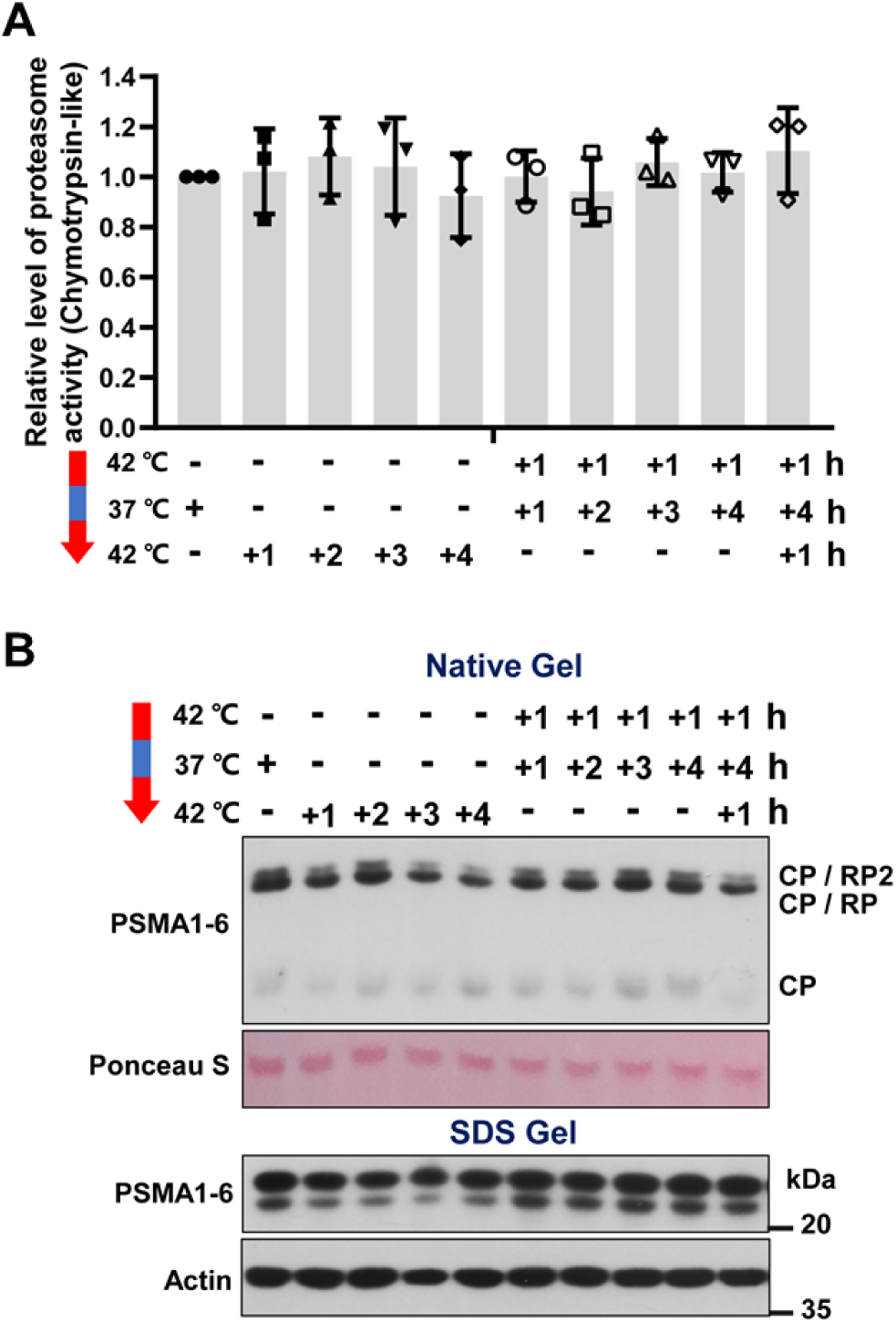
Effect of heat shock and recovery on proteasome activity, abundance, and assembly. **A.** The levels of chymotrypsin-like proteasome activity were determined in lysates from naïve MCF7 cells or MCF7^aTT^ cells rendered thermoresistant through the indicated temperature shift scheme (mean activity +/− SD, n = 3). **B.** Native PAGE of MCF7/MCF7^aTT^ cells kept under the indicated conditions followed by immunoblotting with an antibody recognizing 20S subunits (PSMA1-6, top panel). Ponceau S staining of the blotting membrane is shown to document gel loading. The bottom panel shows a conventional immunoblot of the same lysates with actin as a reference.

**Fig. EV3.**
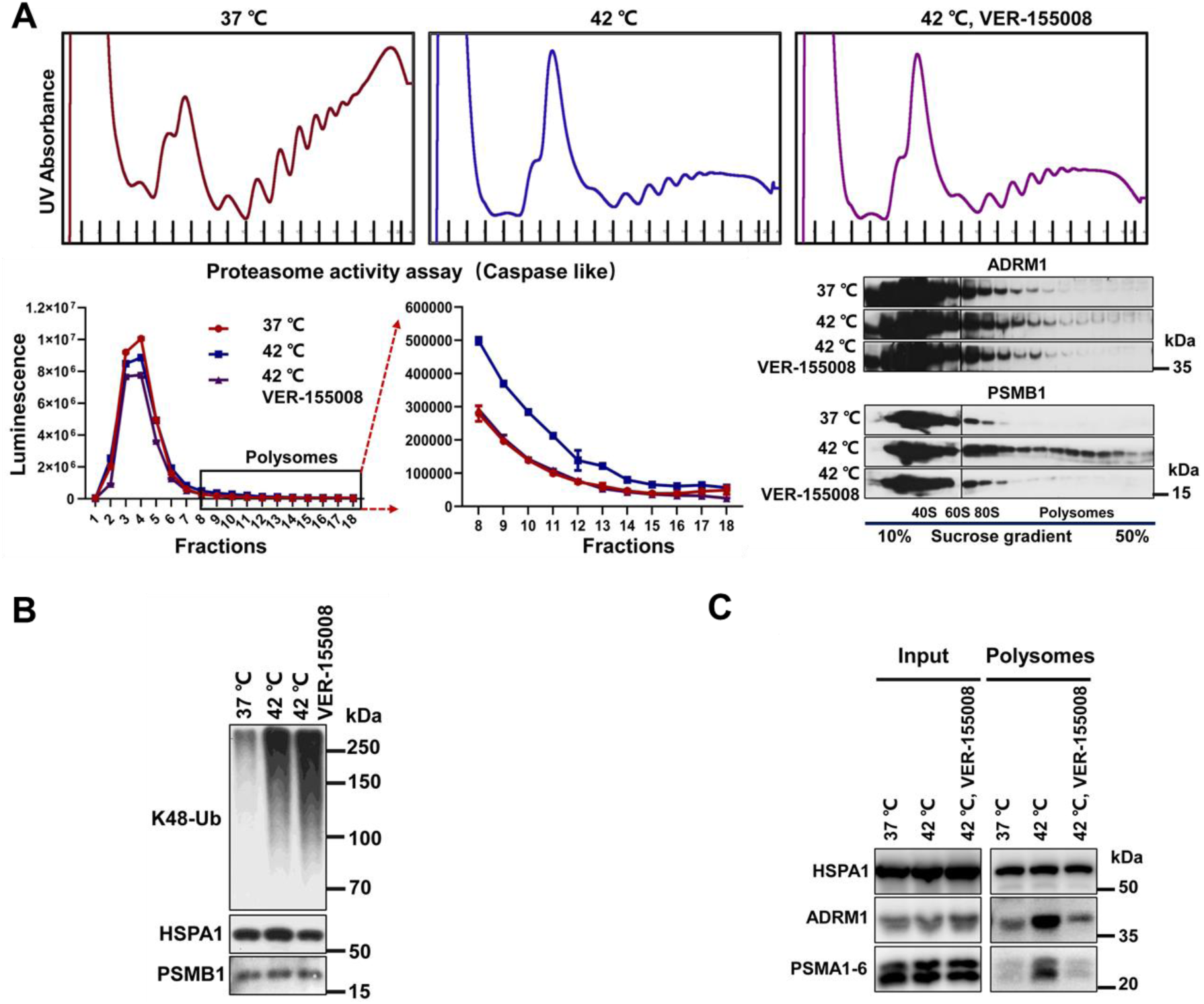
Effect of heat shock and HSP70 inhibition on proteasome recruitment to ribosomes in MCF7 cells. **A.** Heat shock naïve MCF7 cells were exposed to heat shock (42 °C, 1h) or heat shock and HSP70 inhibitor VER-155008 (50 μM, 1h), and resulting cell lysate was separated by sucrose density gradient centrifugation (UV traces, top panel). Caspase-like proteasome activity was assessed in the sucrose density gradient fractions using an in vitro assay. Results are plotted in a line graph. The graph on the right shows a zoom-in on fractions 8 and higher. Fractions were examined by immunoblotting for the elution profiles of the indicated proteasome subunits (right panel). Black vertical lines indicate splicing of lanes that were run on different gels due to limitations in lane capacity. **B.** The unfractionated lysates described in A. were assayed by immunoblotting for the levels of K48-linked polyubiquitin and the indicated proteins. **C.** Lysate of MCF7 cells maintained under the indicated conditions were obtained and polysomes were isolated by centrifugation through a 30% sucrose cushion. Whole cell lysates and the polysomal pellets were assayed by immunoblotting for the levels of HSPA1 and proteasome subunits.

**Fig. EV4.**
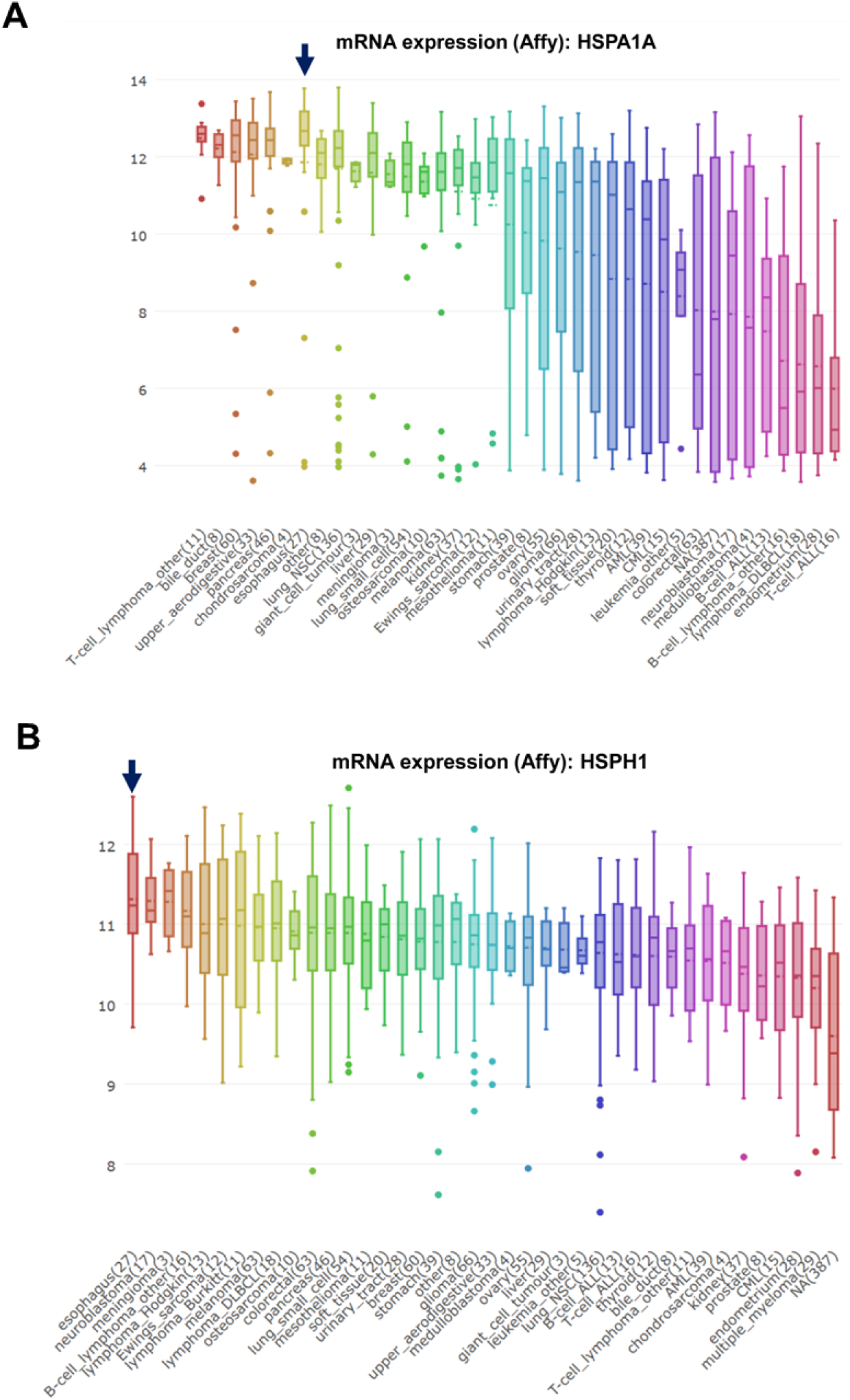
Relative expression of mRNAs encoding HSPA1 and HSPH1 in human cancer cell lines. **A., B.** mRNA expression data were extracted from the MiPanda database (www.mipanda.org, (Niknafs *et al*, 2018).

**Fig. EV5.**
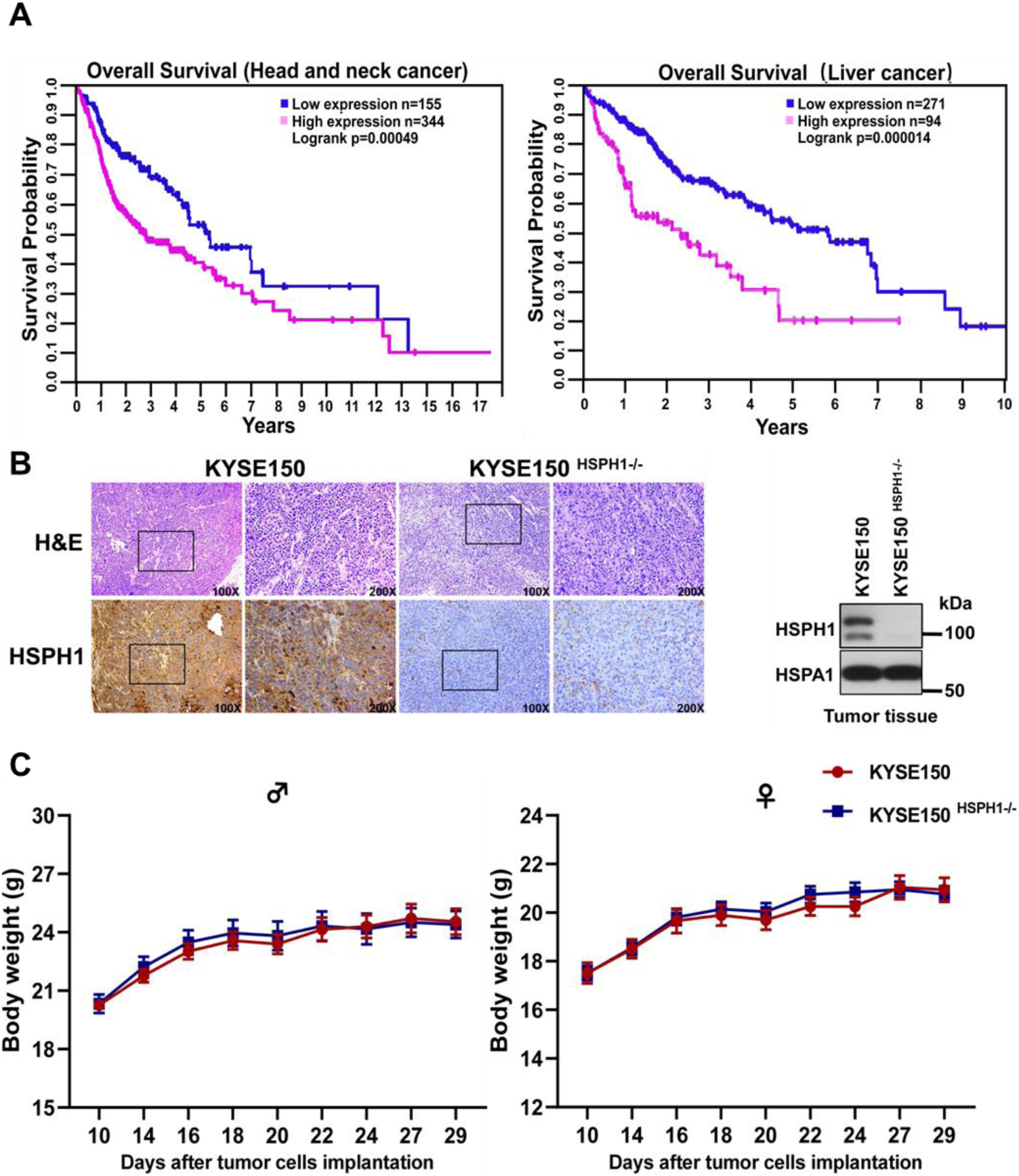
**A.** Kaplan-Meier plots showing the correlation between HSPH1 mRNA expression level and the survival of patients with head and neck or liver cancer. The expression data was obtained from and visualized with KM Plotter (www.kmplot.com, (Nagy *et al*, 2018) using the Pan Cancer algorithm at default settings. **B.** IHC staining of HSPH1 in a representative matched pair of tumors derived from KYSE150 and KYSE150^HSPH1-/-^ cells (left panel). Total protein extracted from one set of matched tumors were assayed for the levels of HSPH1 by immunoblotting (right panel). HSPA1 levels are shown as a reference. **C.** Weights of the animals used for KYSE xenograft studies (means +/− SEM, n = 10).

